# Long-read sequence analysis of MMEJ-mediated CRISPR genome editing reveals complex on-target vector insertions that may escape standard PCR-based quality control

**DOI:** 10.1101/2023.03.03.531065

**Authors:** Yuki Higashitani, Kyoji Horie

**Affiliations:** Department of Physiology II, Nara Medical University, Kashihara, Nara 634-8521, Japan

## Abstract

CRISPR genome editing is a powerful tool for elucidating biological functions. To modify the genome as intended, it is essential to understand the various modes of recombination that can occur. In this study, we report unexpected complex vector insertions that were identified during the generation of conditional alleles by CRISPR editing using microhomology-mediated end joining (MMEJ). The targeting vector contained two loxP sequences and flanking 40-bp microhomologies. The genomic regions corresponding to the loxP sequences were cleaved with Cas9 in mouse embryonic stem cells. PCR screening for targeted recombination revealed a high frequency of bands of a larger size than expected. Nanopore sequencing of these bands revealed complex vector insertions mediated not only by MMEJ but also by non-homologous end joining and homologous recombination in at least 17% of the clones. A new band appeared upon improving the PCR conditions, suggesting the presence of unexpectedly modified alleles that escape standard PCR screening. This prompted us to characterize the recombination of each allele of the genome-edited clones using heterozygous single nucleotide polymorphisms, leading to confirmation of the presence of homozygous alleles. Our study indicates that careful quality control of genome-edited clones is needed to exclude complex, unexpected, on-target vector insertion.

## INTRODUCTION

In recent years, genome modification technology has progressed greatly with the advent of the CRISPR system (1). However, it is always possible for unintended recombination to occur in CRISPR genome editing. Therefore, quality control of genome-edited clones is important to ensure that the intended recombination in the clones is not accompanied by unintended recombination. Unintended recombination may occur at the target site of recombination or in genomic regions distant from the target site. The latter type of recombination often occurs in genomic sequences similar to the target sequence due to the partial annealing of guide RNAs (gRNAs). Such recombination is referred to as “off-target” effects and various methods have been developed to detect it (2,3). Regarding the former unintended “on-target” recombination, large deletions across the target site have been reported (4-7). Typically, recombinants of interest are screened by PCR. Therefore, large genomic deletions that accompany the loss of PCR primer binding sites may go unnoticed. Recently, other types of unintended on-target recombination have been reported in which gRNA vectors or mitochondrial DNA are inserted at the target genomic cleavage site (8). Although the PCR primer binding sites were preserved in these cases, the insertion of the unintended exogenous sequence went unnoticed in the initial PCR screening because the PCR amplicon size was above the detection limit. Such unintended on-target recombination needs to be carefully examined, especially when biallelic genome editing is intended, as loss of the wild-type (wt) allele as verified by PCR analysis may be caused not by homozygous genome editing but by the combination of intended recombination in one allele and unintended recombination in the other allele.

In the present study, we report on unintended, complex, on-target vector insertion patterns that we identified in the process of generating conditional alleles for the Cre/loxP system in cultured cells. To generate a conditional allele, the loxP sequence must be inserted on both sides of the target genomic region. Furthermore, to perform phenotypic analysis in cultured cells, both alleles must be modified. Thus, a total of four locations in the genome must be modified. Different repair pathways, that is, homologous recombination (HR), non-homologous end joining (NHEJ), and microhomology-mediated end joining (MMEJ), can be employed to introduce foreign sequences into the genome. HR is the most widely used pathway and requires a homology arm of 500–1,000 bp in the targeting vector (9). The HR pathway is reported to be active mainly from the late S to the G2 phase of the cell cycle (10). In contrast, the NHEJ pathway is active throughout the cell cycle (10). Therefore, the NHEJ-based method allows for targeted insertion in cell types in which the HR method is inefficient, such as neurons, although the junction sequence at the recombination site is difficult to control compared to that in HR (11). More recently, a method called the PITCh (Precise Integration into Target Chromosome) system using MMEJ was developed (12,13). This method utilizes microhomology arms as short as 10–40 bases in length for recombination. The MMEJ pathway is reported to be active mainly from the M to the early S phase (14), allowing for a different mode of recombination compared to HR. We employed the PITCh method in the present study because the insertion of the 34-base loxP sequence into the genome can be easily detected by PCR due to the short length of the homology arm of the targeting vector. Unexpectedly, PCR screening revealed bands of unintended sizes that were longer than expected at a high frequency. Analysis of these bands with Nanopore long-read sequencing revealed complex vector insertions involving not only MMEJ but also HR and NHEJ. The size of the inserted vector sequence could be beyond the detection limit of the standard PCR-based quality control method for genome-edited clones. We propose that the unintended on-target recombination detected in this study needs to be carefully excluded in the quality control stage of genome editing.

## MATERIALS AND METHODS

### Cell lines and cell culture

The GNL35REC4dn7 mouse ESC line (unpublished), a derivative of the KY1.1 mouse ESC line (15), was used in this study. KY1.1 is a hybrid of the C57BL/6J and 129S6/SvEvTac mouse genetic backgrounds; therefore, heterozygous SNPs can be used to distinguish alleles. The GNL35REC4dn7 ESC line contains the Nano-lantern fluorescent protein gene (16) and the ERT2-iCre-ERT2 cassette (17) at the Gm13227 and Rosa26 loci (18), respectively. These genetic modifications are irrelevant to the purpose of the present study. The ESCs were cultured in KnockOut DMEM (Thermo Fisher Scientific; 10829018) supplemented with 20% fetal bovine serum (FBS), nonessential amino acids (Thermo Fisher Scientific; 11140050), 0.1 mM 2-mercaptoethanol, 1,000 U/ml leukemia inhibitory factor (LIF; Millipore; ESG1107), 1 μM of the MEK inhibitor PD0325901 (Axon Medchem; 1408) and 3 μM of the GSK3 inhibitor CHIR99021 (Axon Medchem; 1386). By adding PD0325901 and CHIR99021, called 2i, to the culture medium, ESCs could be cultured in an undifferentiated state in gelatin-coated dishes without feeder cells. The absence of feeder cells allowed for avoiding the contamination of wt cells in the PCR analysis.

### Construction of the Cas9/gRNA expression vectors

Both Cas9 and gRNAs were expressed using the pX330 vector (19). Complementary oligonucleotides corresponding to the PITCh-gRNA, Klf2-upstream gRNA (gRNA-1), and Klf2-downstream gRNA (gRNA-2) were annealed and cloned into the BbsI site of pX330, resulting in the generation of pX330-PITCh, pX330-gRNA-1 and pX330-gRNA-2 respectively. The oligonucleotides used for vector construction are listed in Supplementary Table 1.

### Construction of the Klf2 targeting vectors

The Klf2 targeting vector was constructed as follows using the primers listed in Supplementary Table 1 and the full nucleotide sequence of the targeting vector is shown in Supplementary Fig. 1. pPGKneo-F2F (gift from Dr. K. Yusa) containing the FRT-flanked neo-selection cassette was digested with SacII and NotI adjacent to the neo cassette. In parallel, genomic fragments of the mouse Klf2 gene from the promoter region to intron 1 and from intron 1 to intron 2 were amplified by PCR using the primer pairs Klf2-5HR-F1 and Klf2-5HR-R1 and Klf2-5HR-F2 and Klf2-5HR-R2, respectively. The 5′ ends of Klf2-5HR-R1 and Klf2-5HR-F2 are complementary to each other and contain the loxP sequence. The 5′ ends of Klf2-5HR-F1 and Klf2-5HR-R2 are complementary to the pPGKneo-F2F vector sequence adjacent to the SacII and NotI digestion sites, respectively. Therefore, the two PCR fragments could be assembled and cloned into the SacII-NotI site of pPGKneo-F2F using the In-Fusion Cloning HD Kit (Takara; 639648), resulting in the generation of pPGKneo-F2F-Klf2-5HR. The Klf2 genomic fragment from intron 2 to the exon 3 untranslated region was synthesized (GenScript) and cloned into the HindIII-SbfI site of pPGKneo-F2F-Klf2-5HR located at the opposite side of the NotI site with respect to the neo cassette, resulting in the generation of pPGKneo-F2F-Klf2-5HR-3HR-mod. A loxP site was introduced in the synthetic Klf2 fragment such that it was located adjacent to the FRT-flanked neo cassette of pPGKneo-F2F-Klf2-5HR-3HR-mod. To generate the PITCh-targeting vector, a DNA fragment was synthesized containing a PITCh-gRNA target site, 5′ microhomology sequence, loxP site, BglII and AscI site, 3′ microhomology sequence, and PITCh-gRNA target site (12) in this order and cloned into the pUC57 vector, resulting in the generation of pPITCh-bk. The BglII-AscI fragment of pPGKneo-F2F-Klf2-5HR-3HR-mod containing the Klf2 intron1-intron 2 fragment, FRT-flanked neo cassette, and a loxP site was cloned into the BglII-AscI site of pPITCh-bk, resulting in the production of the PITCh targeting vector pKlf2-cKO-PITCh.

### Transfection, colony isolation, and DNA preparation

On day 0, ESCs (2.5 × 10^5^) were mixed with 1.25 μg of pKlf2-cKO-PITCh, 0.42 μg of each Cas9/gRNA vector (pX330-PITCh, pX330-gRNA-1, and pX330-gRNA-2), and 5 μl of TransFast (Promega; E2431) in a serum-containing medium in a total volume of 500 μl and plated into one well of a 24-well plate. After 1 h, 1 ml of medium was added to the well, and the medium was replaced with fresh medium 3 h after transfection. On day 1, ESCs were passaged to a 6-cm dish and selected using 250 μg/ml of G418. On day 9, ESC colonies were picked and divided into lysate preparations for PCR analysis and cell cultures for re-cloning and preparing frozen cell stocks. ESC lysates were prepared by resuspending ESC pellets in 20 μl of water and heating at 95 °C for 10 min, followed by Proteinase K digestion (400 μg/ml) at 56 °C for 60 min and heat inactivation of Proteinase K at 95 °C for 10 min. For a detailed analysis of the inserted sequence, some ESC clones were plated sparsely in culture dishes to obtain single-cell–derived colonies. After picking colonies, ESCs were expanded and genomic DNA was purified by the conventional method of phenol/chloroform extraction and ethanol precipitation (20).

### PCR analysis

Targeted clones were screened by PCR using KOD FX polymerase (TOYOBO; KFX-101) with the primers shown in Supplementary Table 1. For the first screening of the targeted clones (Fig. 2B), ESC lysate was used as a template. For upstream and downstream recombination (primer pairs I and III in Fig. 2), the PCR conditions were as follows: one cycle at 94 °C for 2 min, followed by 35 cycles of denaturation at 98 °C for 10 s, annealing at 55 °C for 30 s, and extension at 68 °C for 30 s, with a final extension step at 68 °C for 1 min. For the amplification of full-length insertion sequences (primer pair II in Fig. 2), the extension time of the PCR cycle and the final extension time after the PCR cycle were set as 3 min. When analyzing re-cloned cells (Fig. 2C), genomic DNA purified via phenol/chloroform extraction and ethanol precipitation was used as a template. The PCR conditions were the same as for the lysate analysis except that the extension time of the PCR cycle was set as 8 min and the final extension time after the PCR cycle was set as 5 min to allow for the detection of unexpectedly long insertion sequences.

**Figure 1.**
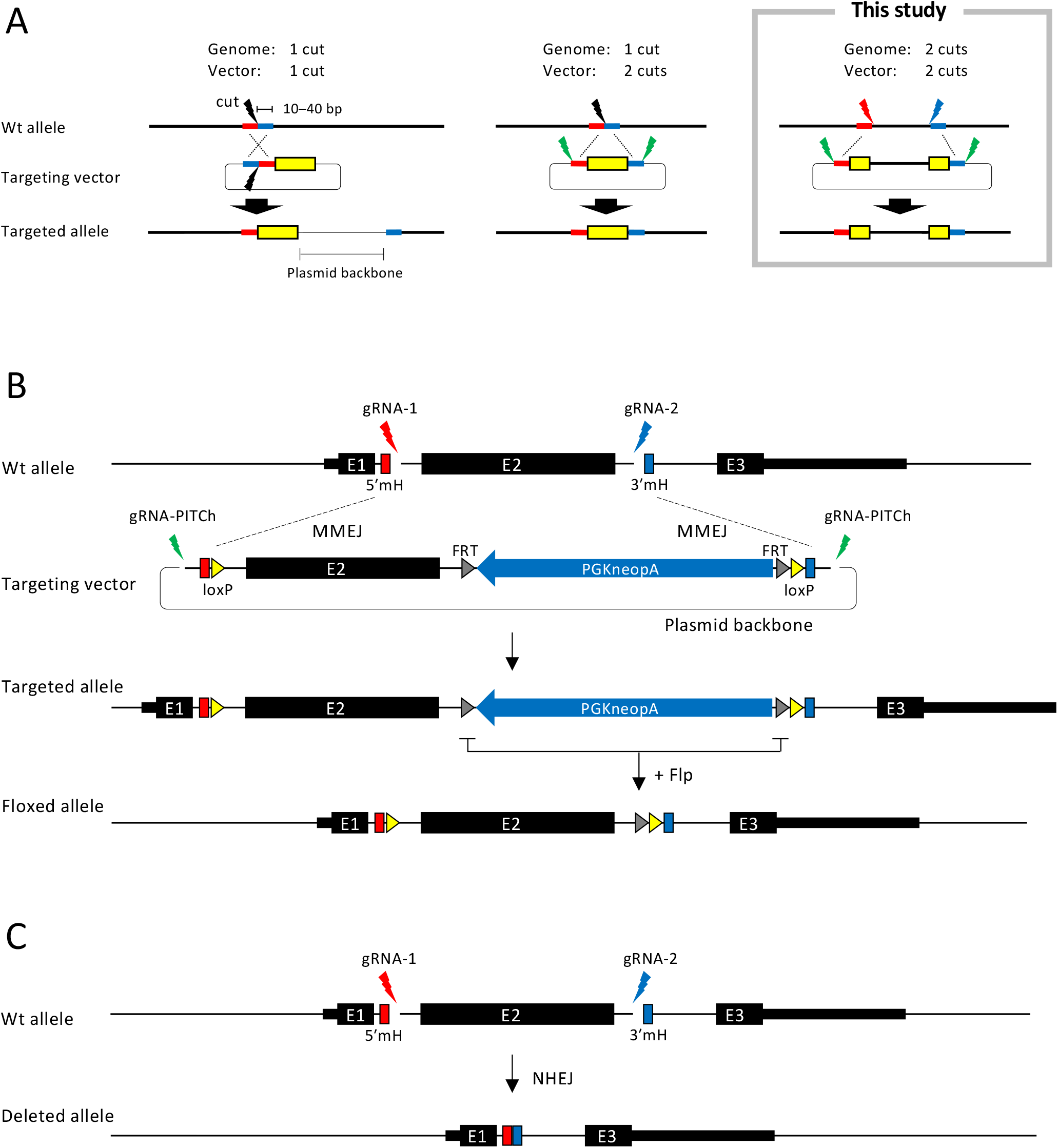
Overview of the MMEJ-mediated targeting strategy. (A) The MMEJ-mediated genome-editing system PITCh, classified by the number of cutting sites in the genome and the targeting vector. Thick red and blue lines, microhomologies; yellow box, exogenous sequence. (B) Generation of the conditional allele at the mouse Klf2 locus. E, exon; 5′mH, 5′ microhomology; 3′mH, 3′ microhomology; PGK, phosphoglycerate kinase-1 promoter; neo, neomycin-resistance gene; pA, polyadenylation signal. (C) Deleted allele generated when the targeting vector was not involved in recombination. The size of each element is not to exact scale due to space limitations.

**Figure 2.**
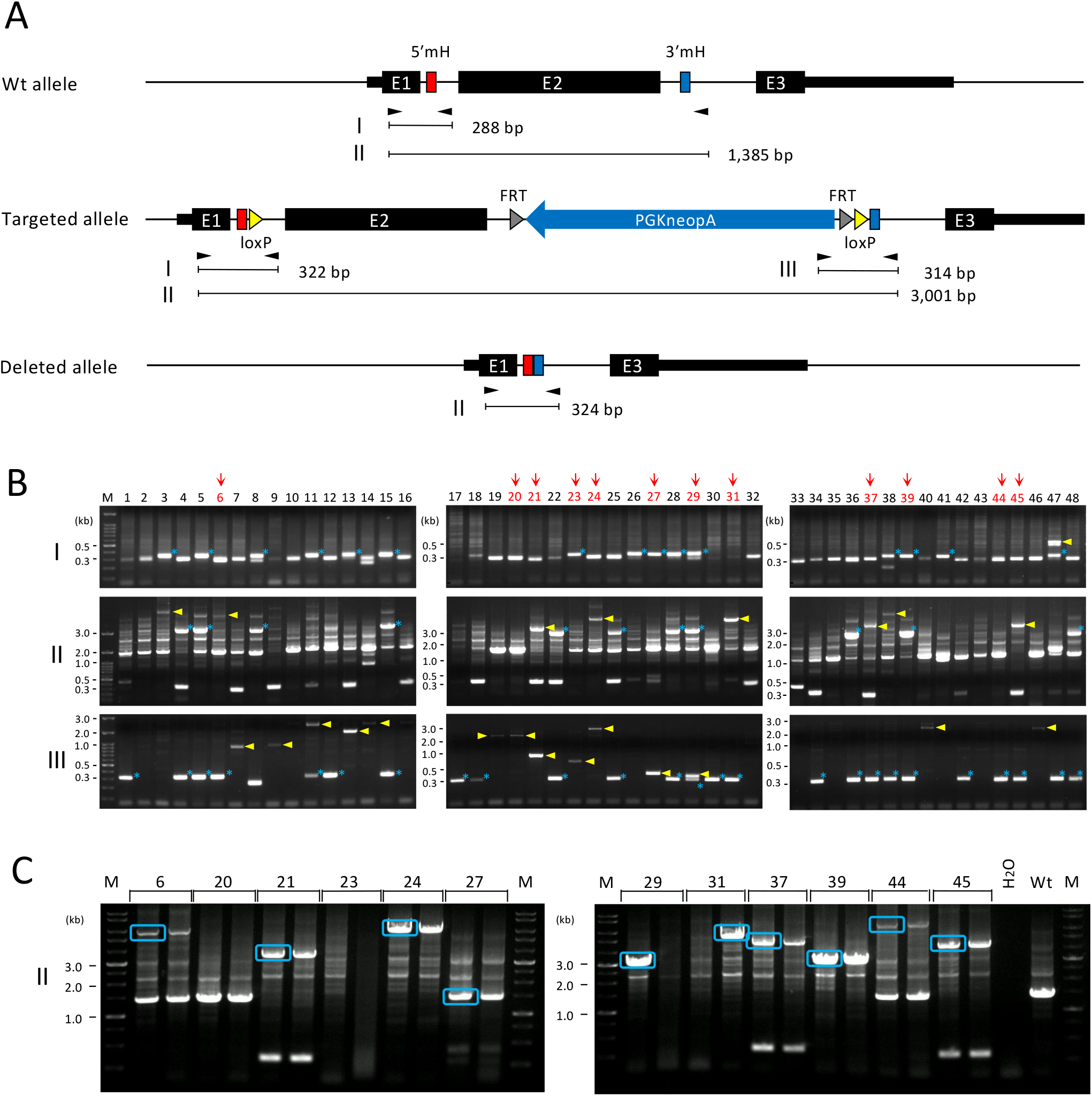
PCR screening for targeted clones. (A) Genomic structure of the wild-type, targeted, and deleted alleles. Black arrowheads indicate PCR primers. Other abbreviations are the same as in Fig. 1B. (B) Results of the PCR screening. The results in panels I, II, and III were obtained using the PCR primer pairs I, II, and III shown in (A), respectively. Bands close to the expected size are indicated by asterisks and bands longer than expected are indicated by arrowheads. Clones marked with arrows were re-cloned and analyzed as shown in (C). M, DNA size marker. (C) PCR analysis after re-cloning. Bands surrounded by rectangles were purified and further analyzed by long-read sequencing.

### Nanopore sequencing

PCR products were purified using the QIAquick Gel Extraction Kit (Qiagen; 28704) and cloned into a plasmid using the Zero Blunt TOPO PCR Cloning Kit (Thermo Fisher Scientific; 450245). Plasmid DNA was purified using the QIAprep Spin Miniprep Kit (Qiagen; 27106), digested with BsrGI, and purified with the QIAquick PCR Purification Kit (Qiagen; 28104). Long-read sequencing was performed on a MinION sequencer (Oxford Nanopore Technologies) using the 1D ligation sequencing kit (Oxford Nanopore Technologies; SQK-LSK109) and native barcoding kit (Oxford Nanopore Technologies; EXP-NBD103). Raw sequence reads were assembled using Flye version 2.6 (21).

### Sanger sequencing

PCR products were purified with either the QIAquick PCR Purification Kit (Qiagen; 28104) or the QIAquick Gel Extraction Kit (Qiagen; 28704) and directly sequenced using the BigDye Terminator v3.1 Cycle Sequencing Kit (Thermo Fisher Scientific; 4337455). All Sanger sequencing was conducted on PCR products, not on the plasmid DNA with the PCR-amplified fragment inserted, to avoid any bias caused by PCR-induced mutations.

## RESULTS

### Overview of the targeting strategy

Fig. 1A presents an overview of the MMEJ-mediated genome-editing system PITCh (12,13). The full-length sequence of the targeting vector is shown in Supplementary Fig. 1. The PITCh method can be classified into three types according to the number of cutting sites in the genome and the targeting vector. Cutting the genome and targeting vector at one site is straightforward but the plasmid backbone of the targeting vector is incorporated into the genome (Fig. 1A, left). Cutting at one site in the genome and two sites in the targeting vector prevents the incorporation of the plasmid backbone into the genome (Fig. 1A, middle). In the present study, we cut at two sites in both the genome and the targeting vector (Fig. 1A, right). This strategy allows for the insertion of exogenous sequences at two locations in the genome.

Fig. 1B illustrates the strategy for the generation of conditional alleles. We planned to delete exon 2 of the mouse Klf2 gene using the Cre/loxP system in mouse embryonic stem cells (ESCs). We expressed gRNAs as single-guide RNAs (sgRNAs) using the Cas9/gRNA expression vector pX330 (19). LoxP sites were inserted into the targeting vector exactly at the cutting sites of the Cas9/gRNA complex. Forty base pairs of microhomologies were placed upstream and downstream of the loxP sites in the targeting vector. The gRNA target site, which is described in the original report on the PITCh system (12), was inserted outside the microhomologies of the targeting vector (Supplementary Fig. 1). The targeting vector contained the neomycin-resistance gene (neo) cassette to enrich the targeted events by G418 selection.

Mouse ESCs were transfected with the targeting vector and the three gRNAs (Fig. 1B). Successful cleavage of the gRNA target sites would expose microhomologies in the genome and the targeting vector that would be utilized for MMEJ-assisted recombination. Targeted clones were screened by PCR as detailed in the next section. The neo cassette could be excised by the Flp/FRT system after isolating targeted clones. The entire region of exon 2 between the two gRNA target sites could be deleted by NHEJ in case the targeting vector was not involved in recombination (Fig. 1C).

### PCR screening for the targeted clones

Fig. 2 shows the results of the PCR screening for the targeted clones. Primer pair I detects upstream recombination: a 288-bp band is expected from the wt allele and a 322-bp band from the expected recombination (Fig. 2A). Primer pair II amplifies the target sites spanning the vector insertion: a 1,385-bp band is expected from the wt allele, a 3,001-bp band from the predicted recombination, and a 324-bp band from the deletion between the two cleavage sites (Fig. 2A). Primer pair III detects downstream recombination: a 314-bp band is expected from the intended recombination (Fig. 2A).

We screened 48 clones in total (Fig. 2B). A band of around 0.3 kb was observed in some clones with primer pair II, indicating a deletion between the two cutting sites (Fig. 2A). Using primer pairs I, II, and III, bands expected from the targeted allele were detected in 15, 11, and 25 clones, respectively (indicated by asterisks in Fig. 2B). However, only five clones showed the expected bands with all primer pairs (clones 5, 15, 28, 29, and 39 in Fig. 2B), indicating that unexpected vector insertion patterns frequently occurred. Longer-than-expected bands (depicted by arrowheads in Fig. 2B) were observed in 21 clones, further indicating that the vector insertion pattern was different than expected. Theoretically, an insertion detected with primer pair III can be detected with primer pair II if the upstream primer is present and the amplification efficiency is sufficient. However, out of the 14 clones that showed longer-than-expected bands with primer pair III, only two clones (clones 21 and 24) showed corresponding amplification with primer pair II, suggesting the possibility that most of the vector insertions were beyond the detection limit with primer pair II.

Based on these results, we considered that the nature of the unpredicted vector insertions needs to be elucidated to accurately assess the quality of the genome-edited clones. To obtain clues regarding the mechanism of the unpredicted recombination, we aimed to determine the sequence of the entire region of unexpected insertions. We selected the following clones, indicated by arrows in Fig. 2B: first, clones that showed a larger-than-expected band size for at least one of the primer pairs (clones 6, 21, 24, 31, 37, and 45) and second, clones with no PCR amplification other than the presence of a wt-sized band for primer pair II, although amplification for either primer pair I or III strongly suggested that vector insertion occurred at the target site (clones 20, 23, 27, and 44). Third, we included two clones showing the expected band size for all primer pairs (clones 29 and 39). Clone 39 showed a very strong 3-kb band for primer pair II accompanied by a very faint 1.4-kb wt-sized band, suggesting that biallelic recombination occurred but a small number of wt allele–containing cells were contaminated. In contrast, clone 29 showed non-negligible amplification of a 1.4-kb wt-sized band for primer pair II in addition to the expected 3-kb band. Clone 29 also showed two bands with different intensities for primer pairs I and III, strongly suggesting the possibility that two clones were fused during the colony formation of G418-resistant ESCs. This result indicates that re-cloning is needed to accurately analyze the recombination event.

Therefore, we performed the re-cloning of each clone by plating cells sparsely in culture dishes and isolating single-cell–derived colonies (Fig. 2C). For the PCR analysis in Fig. 2B, cell lysates were used as templates to allow for rapid screening (see MATERIALS AND METHODS for details of the procedure). On the other hand, to increase the detection sensitivity of PCR, the genomic DNA of the re-cloned cells was prepared to high purity by phenol/chloroform extraction and ethanol precipitation. For each clone, two subclones were isolated and subjected to PCR analysis (Fig. 2C). In the PCR screening in Fig. 2, the elongation time was 3 min. On the other hand, the elongation time was set as 8 min in the analysis of the re-cloned cells so that long insertion sequences could be amplified. Many clones showed the same pattern in the two subclones (Fig. 2C), in which case we used one of the subclones for further analysis. In some cases, only one of the subclones showed a band or both subclones did not show any bands (Fig. 2C), suggesting that more than one clone was mixed before re-cloning. Only one subclone showed a PCR band for clones 29 and 31, indicating that these clones comprised a mixture of more than one clone. The 1.4-kb wt-sized band that was observed in Fig. 2B disappeared for clone 29, suggesting that this clone may have had the expected vector insertion in both alleles. Importantly, in clone 44, we observed a new band that was larger than predicted (Fig. 2C) and was not visible in the first PCR screening (Fig. 2B). This result demonstrates that an unintended vector insertion pattern can escape detection by the standard PCR screening procedure.

### Long-read sequence analysis of vector insertions

Fig. 3A illustrates the sequencing strategy. First, the PCR band was cloned into a plasmid. Since the PCR band was larger than expected, it is possible that repeat sequences were present in the PCR products, such as the insertion of two vector copies. Therefore, we used the MinION long-read sequencer, which is capable of analyzing repetitive sequences. The consensus sequences of the read sequences were obtained using an assembly tool, Flye (21) (Supplementary Fig. 2). The insertion pattern of the vector was then estimated by comparing the consensus sequences with the vector and genomic sequences. In MinION sequencing, nucleotide sequence errors occur at a high frequency so care must be taken when interpreting the consensus sequence. Therefore, the sequences of the recombination junctions between the genome and the vector or between two vectors, which were estimated by MinION, were also determined by direct Sanger sequencing of the PCR products shown in Fig. 2C. All the target sequences of the gRNAs were also determined by the Sanger method. Even when a gRNA target site was not involved in the recombination with the targeting vector as shown later, the presence of the mutation at the gRNA target site demonstrated that the target sequence was cleaved by the Cas9/gRNA complex.

**Figure 3.**
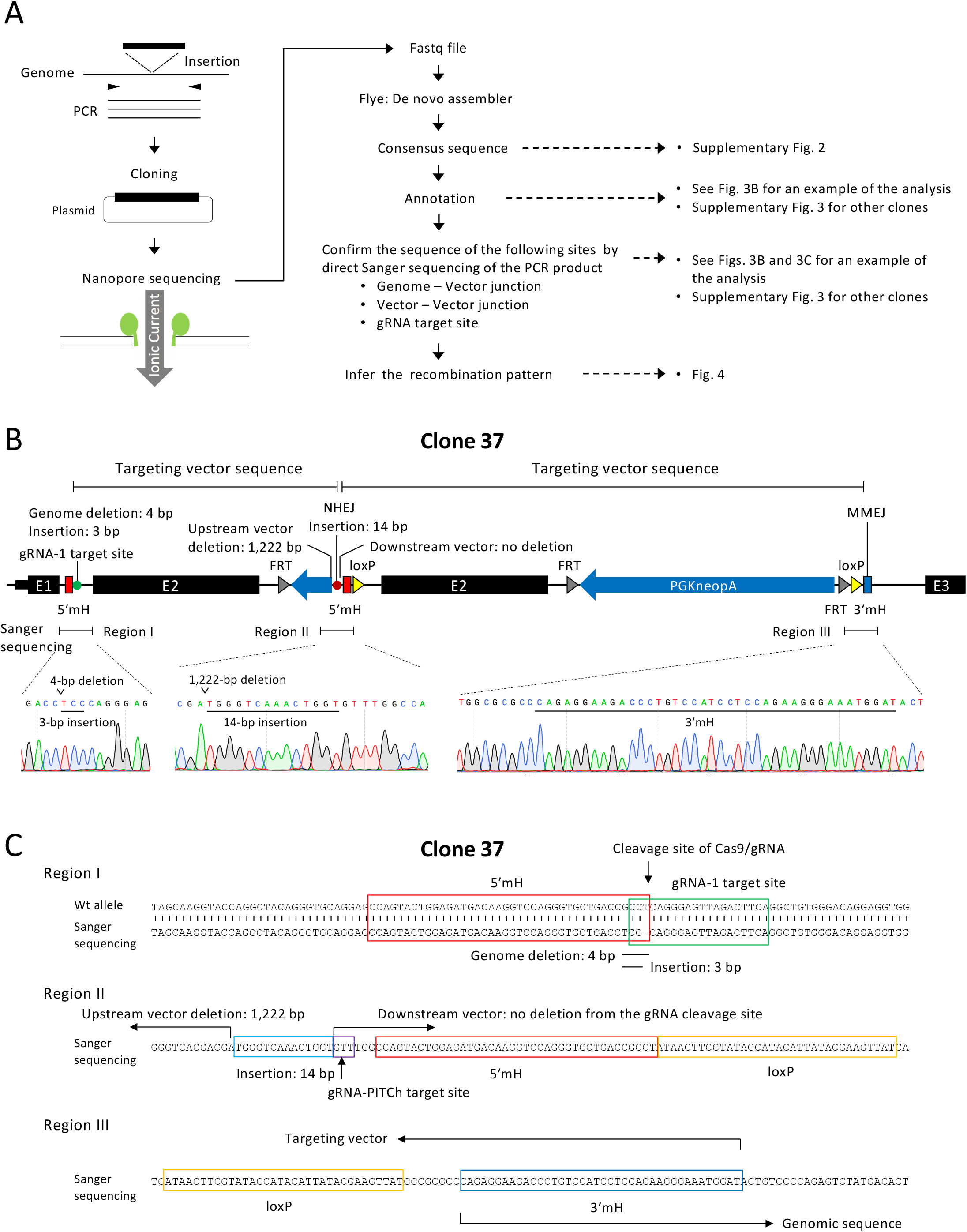
Long-read sequence analysis of vector insertion. (A) Overview of long-read sequence analysis. (B) Example of annotation of a consensus sequence obtained by de novo assembly of long-read sequencing data. Examples of direct sequencing using the Sanger method, performed to confirm the fidelity of the assembly, are also shown. (C) Annotation of the results of Sanger sequencing in regions I, II, and III shown in (B).

Figs. 3B and 3C show an example of the analysis of clone 37 (Fig. 2C), and the recombination pattern inferred from this analysis is shown in Fig. 4F. In this example, two vectors were linked by NHEJ and inserted into the genome. At the vector linkage site, an approximately 1.2-kb sequence was deleted in one of the vectors (Figs. 3B, 3C). The linked vector was inserted into the genome by HR between exon 2 sequences in the upstream region and by the originally envisioned MMEJ in the downstream region (Figs. 3B, 3C, 4F). Although the upstream gRNA target site was not involved in recombination with the vector, a mutation was detected at this site by Sanger sequencing (Figs. 3B, 3C), confirming that cleavage did occur.

**Figure 4.**
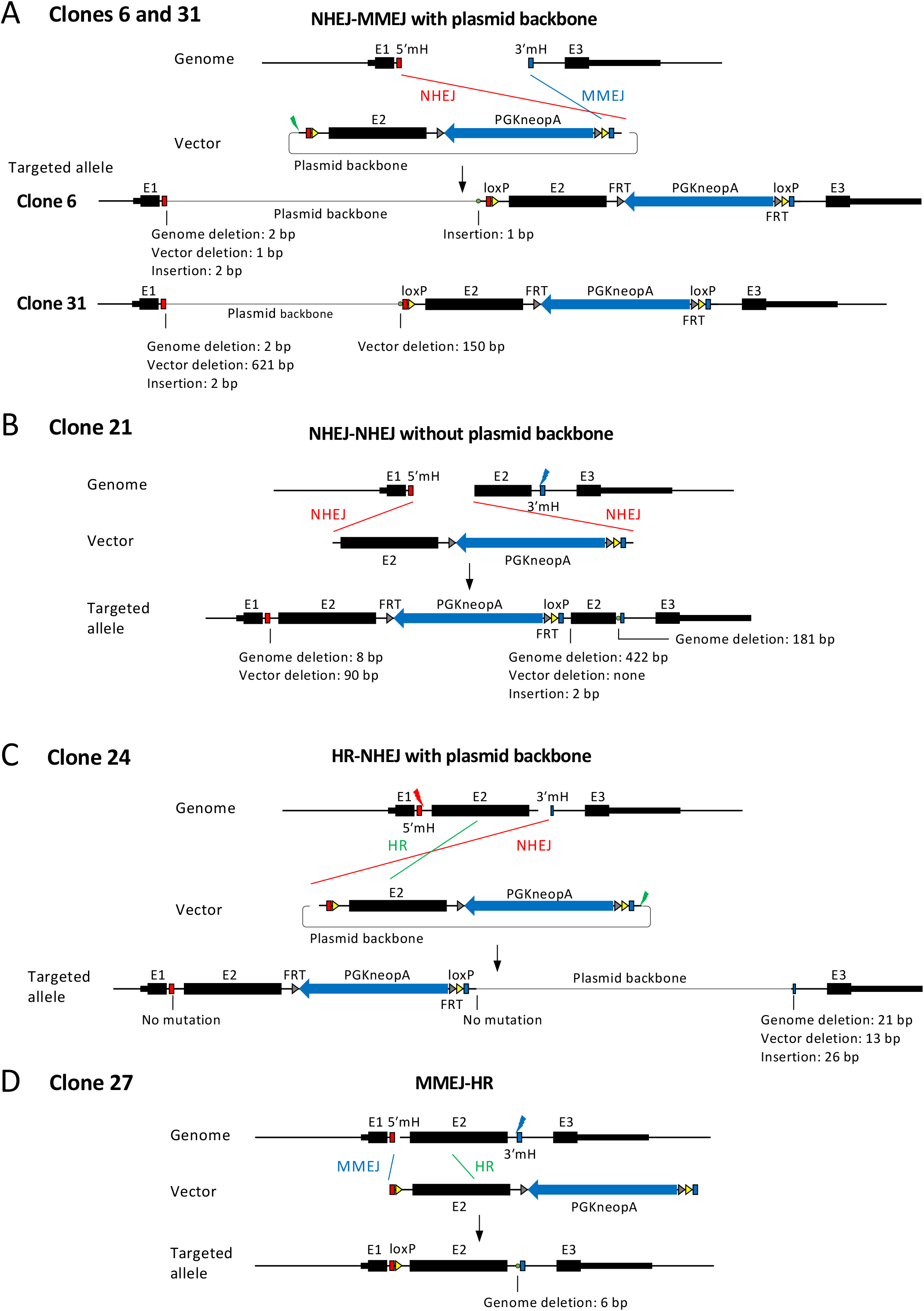
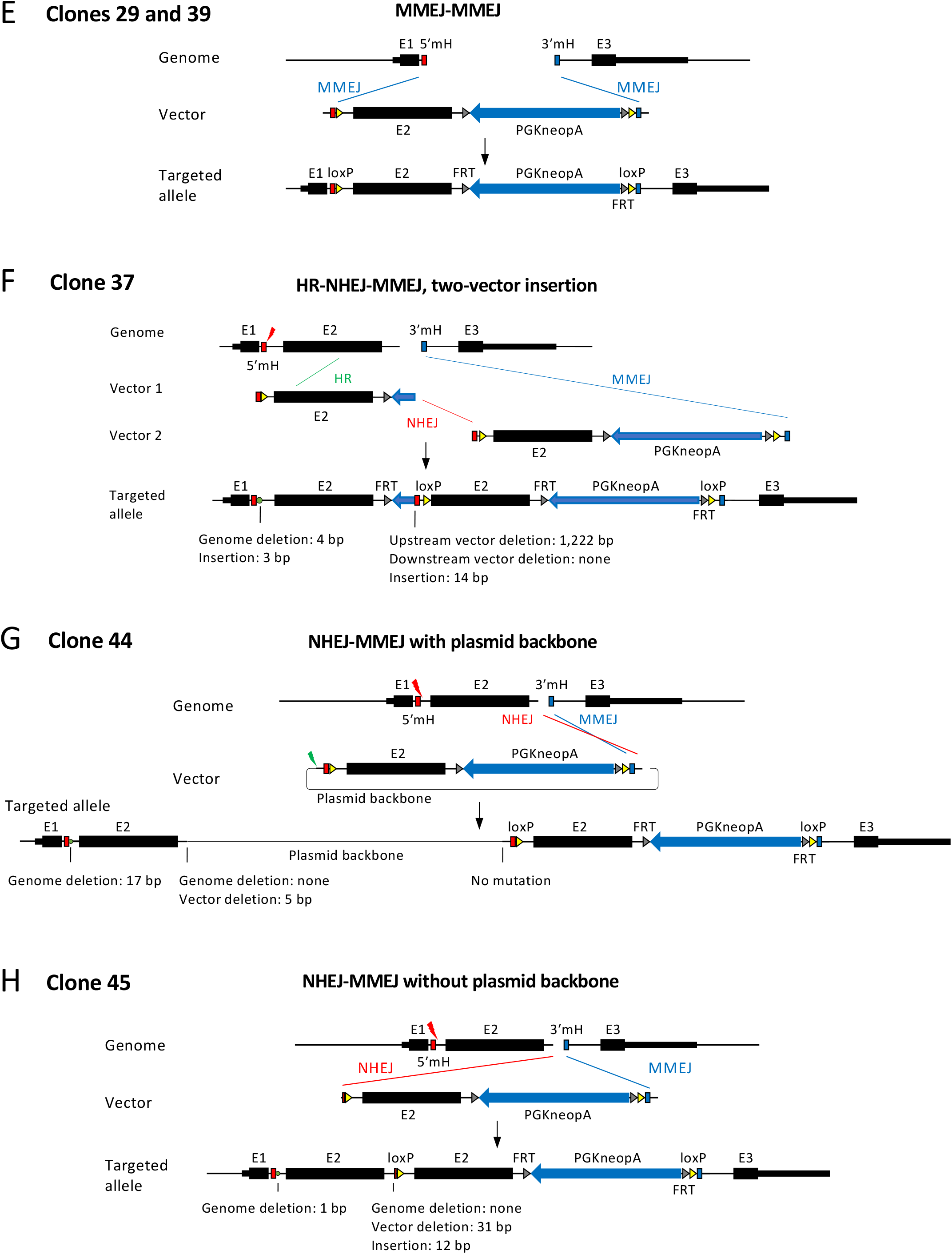
Recombination patterns inferred from long-read sequencing. Each recombination pattern was inferred based on the results shown in Figs. 3B and 3C for clone 37 and Supplementary Fig. 3 for other clones.

The same analysis as in Figs. 3B and 3C was performed for all clones, as detailed in Supplementary Fig. 3. The recombination patterns inferred from the analysis are shown in Fig. 4. In many cases, only one cleavage site of the genome or the vector was involved in recombination (clones 6, 21, 24, 27, 31, 37, 44, and 45). Since a microhomology pair of the genome and the vector must be exposed by a double-strand break for MMEJ to occur (Fig. 1), MMEJ could not occur in these clones in either the upstream or the downstream region. Instead, NHEJ or HR was observed in such regions. As a result, different combinations of repair pathways, that is, NHEJ and MMEJ (clones 6, 31, 44, and 45 in Fig. 4), HR and MMEJ (clone 27 in Fig. 4), HR and NHEJ (clone 24 in Fig. 4), and HR, NHEJ, and MMEJ (clone 37 in Fig. 4), were involved in vector insertions. In one case, NHEJ, not MMEJ, occurred even though microhomology must have been present at the cleavage site (clone 21).

As shown above, recombination at only one cleavage site in the genome or the vector will result in the insertion of an unexpected sequence into the genome. This was found to be one of the reasons for the presence of unexpected PCR bands of larger-than-expected size. If only one cleavage site in the vector is used for recombination, the PCR band size will become considerably larger because of the insertion of the 2.7-kb vector backbone into the genome (clones 6, 24, 31, and 44 in Fig. 4). Furthermore, if only one cleavage site is involved in recombination in both the genome and the vector as in the case of clone 44 (Fig. 4G), the band will be 3.8 kb longer than expected, resulting in a substantial reduction in the detection sensitivity of PCR. In fact, the full-length vector sequence could not be amplified by the initial PCR for clone 44 (Fig. 2B) and was amplified only after increasing the PCR extension time from 3 min to 8 min and using a highly purified genomic DNA template (Fig. 2C). Multiple vector linkage also increases the size of the vector insertion significantly as observed in clone 37 (Fig. 4F).

Interestingly, no selection marker was present at the integration site for clone 27. Since all clones were selected for G418 resistance, another copy of the vector sequence was likely inserted into the genome. The two clones that showed the expected PCR bands for all primer pairs were both found to have undergone MMEJ-mediated recombination as expected (clones 29 and 39).

Taken together, eight clones were identified by long-read sequencing analysis as having unexpected recombination patterns. Since 48 clones were initially screened by PCR, the frequency of unexpected recombination patterns was 17% (eight out of 48). We consider this frequency to be an underestimate because 11 more clones showed longer-than-expected PCR bands (Fig. 2B) but were not analyzed in detail (see DISCUSSION).

### Allelic analysis of recombination using heterozygous single nucleotide polymorphisms (SNPs)

The results shown above indicate that unexpected recombination may generate alleles that can escape detection by PCR. This is especially important when aiming for homozygous genome modification, which was the original aim of this study. Clones 29 and 39 comprised candidates for homozygous genome modification because the expected MMEJ-induced recombination was demonstrated by sequence analysis (Fig. 4E) and no wt alleles were detected by PCR after re-cloning (Fig. 2C). However, the possibility remains that MMEJ-mediated recombination occurred only in one allele, while a large sequence not detectable by PCR was inserted in the other allele. To exclude this possibility, we redesigned the PCR screening. A primer pair was set up to amplify heterozygous SNPs upstream and downstream of the target site (Fig. 5A). Using this primer pair, we reanalyzed clones 29 and 39 and obtained a single band as expected (Fig. 5B, left). Sequencing of both ends of this band revealed the presence of heterozygous SNPs (Fig. 5B, right). This confirmed that the predicted recombination occurred in both alleles of these clones.

**Figure 5.**
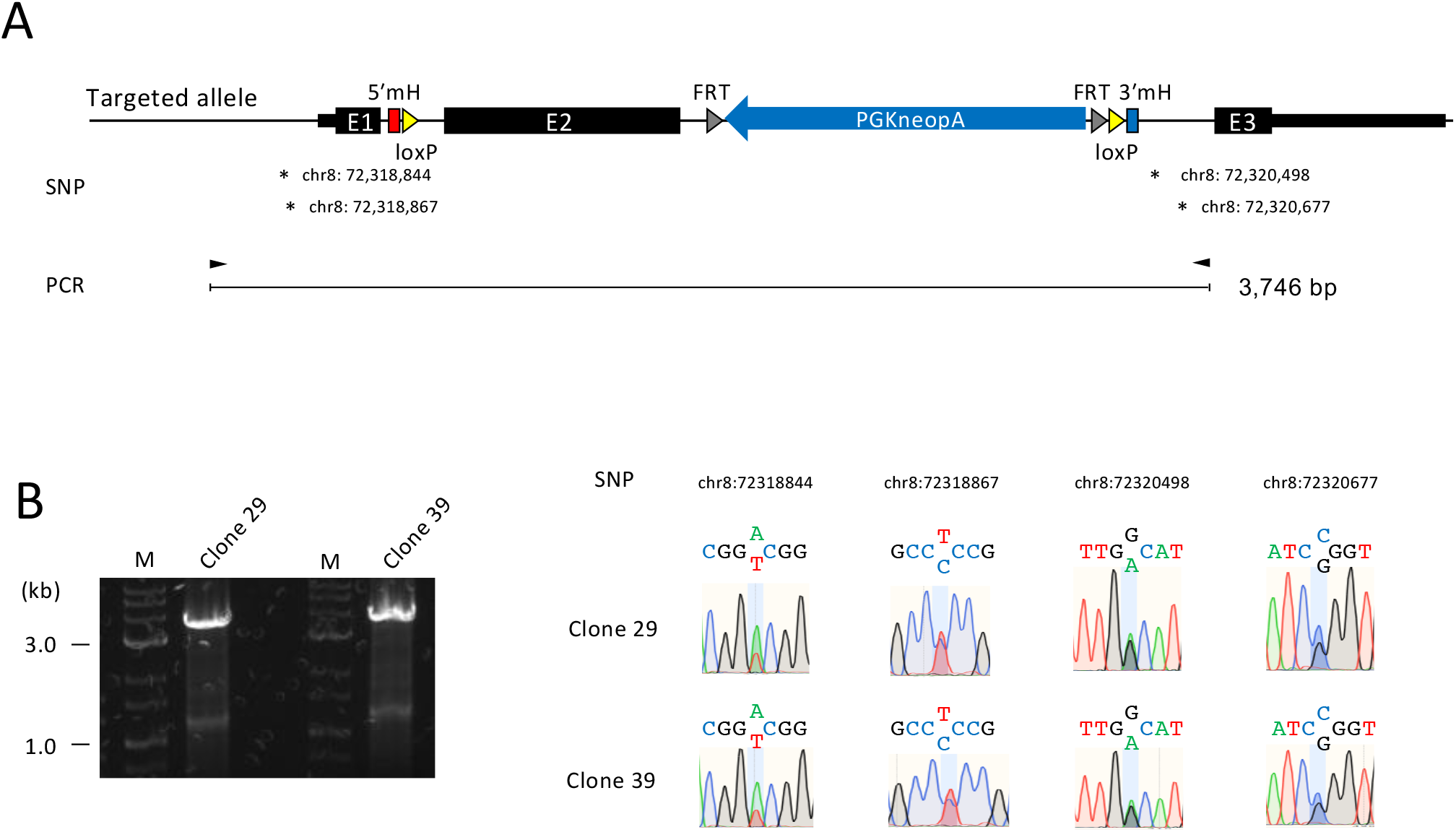
Allelic analysis of recombination using heterozygous SNPs. (A) Structure of the targeted allele of clones 29 and 39. Locations of the heterozygous SNPs to distinguish the C57BL/6J and 129S6/SvEvTac alleles are shown by the genomic coordinates of mm10 genome assembly. (B) PCR product of the full-length insertion sequence (left) and SNP analysis by Sanger sequencing (right). Heterozygosity was detected at all SNPs, indicating homozygously targeted events. M, DNA size marker.

## DISCUSSION

The present results indicate that unexpectedly long exogenous sequences that escape detection by conventional PCR screening can be inserted at the target site in genome editing. It is important to elucidate the mechanism underlying such unexpected recombination to accurately assess genome-edited clones. For example, loss of the wt allele by PCR may be misinterpreted as a homozygously targeted event rather than being attributed to the insertion of long exogenous sequences beyond the detection limit of PCR. To obtain clues for the characterization of such unexpected recombination events, we analyzed clones containing insertions of unexpectedly long exogenous sequences using Nanopore long-read sequencing technology. The analyses revealed the involvement of unintended repair pathways such as NHEJ and HR in vector insertion, resulting in recombination that may be difficult to detect by conventional PCR screening. Based on these findings, we supposed the importance of distinguishing recombination between alleles and redesigned the PCR analysis using heterozygous SNPs (Fig. 5).

In total, we determined the full-length sequences of vector insertions in eight clones out of 48 clones screened by PCR (Figs. 3, 4, Supplementary Figs. 2, 3). Therefore, 17% of the insertions were demonstrated to be unpredicted patterns involving not only MMEJ but also NHEJ and HR. It should be noted, however, that this frequency is an underestimate because an additional 11 clones showed longer-than-expected bands in the initial PCR screening but were not analyzed further (Fig. 2B). If we include these 11 clones, the frequency of unexpected recombination reaches 40%. This observation strongly suggests that great care should be taken in the quality control of genome-edited clones.

To generate conditional alleles at the mouse Klf2 gene locus, we used an MMEJ-mediated genome-editing method called PITCh. HR-mediated methods are more frequently used for the insertion of exogenous sequences. HR uses 500–1,000-bp homologous sequences in the targeting vectors (9), whereas MMEJ uses short homologous sequences of 5 to 40 bp (12,13,22,23). In the present study, we aimed to insert a loxP sequence of only 34 bp. Therefore, we considered the PITCh method, with its short homology arm, to be more suited to detecting the increase in the length of the PCR product after recombination compared to the HR-mediated method using longer homology arms (Fig. 2B). However, we observed a high frequency of unexpected recombination. One possible reason for this is that both the genome and the vector needed to be cleaved by Cas9/gRNA at two locations (Fig. 1B) but were, in reality, not cleaved simultaneously in cultured cells. To our knowledge, there are two reports of the MMEJ-based PITCh method in which both the genome and the vector were cut at two locations. In both studies, the expected recombination was observed at a higher frequency than in our case (22,23). However, both experiments were performed using fertilized mouse eggs. All gRNA target sites would be cleaved seamlessly in fertilized eggs because they can be injected with high concentrations of Cas9, gRNA, and targeting vectors. Therefore, a direct comparison with the present study, which used cultured cells, is difficult. In the present study, only one gRNA target site in either the genome or the vector was involved in the recombination between the genome and the vector in eight clones (clones 6, 21, 24, 27, 31, 37, 44, and 45 in Fig. 4). All gRNA target sites not involved in recombination were mutated in six out of eight clones (Fig. 4), indicating that gRNA cleavage did occur. Perhaps the gRNA target sites were cleaved at different times and, therefore, could not be utilized efficiently for MMEJ-mediated recombination. The cleaved ends of the targeting vector in PITCh would result in NHEJ-mediated vector insertion into the genome when microhomology in the genome was not exposed by a double-strand break. In contrast, in conventional HR-based genome editing, targeting vectors are used in a circular form, which may reduce the frequency of vector insertion via NHEJ. However, a recent report showed that a circular gRNA expression vector and mitochondrial DNA were inserted into genomic cleavage sites during genome editing using CRISPR-Cas9 (8). Therefore, even with HR using a circular targeting vector, the vector could be inserted into the genome in an unexpected manner. A study showed that linearizing the targeting vector improves the efficiency of HR-based genome editing (24). The authors also noted that NHEJ can occur between genomic cleavage sites and linearized vectors but did not determine the entire insertion sequence, as has been done in the present study. Based on the results of our study, we believe that when using linearized vectors in HR-based genome editing, sufficient attention should be paid to unexpected vector insertions that are difficult to detect by PCR.

PCR analysis after re-cloning showed increased amplification of the unexpected bands or the appearance of new bands that were undetectable before re-cloning (Fig. 2C). One possible reason for this is that the extension time was increased from 3 to 8 min. Another factor may have been the use of genomic DNA highly purified by phenol/chloroform extraction and ethanol precipitation as the PCR template after re-cloning, whereas cell lysates were used as the template before re-cloning. However, if the inserted sequence is extremely long, PCR amplification would fail even under improved conditions. For example, three or more targeting vectors could be linked and inserted into the genome, and such recombination would be beyond the detection limit of PCR. In fact, when transgenic mice are created by the microinjection of vector DNA into fertilized eggs, hundreds of copies of tandemly arrayed vector DNAs can be inserted into specific sites in the genome (25), and the same thing may happen in cultured cells. Therefore, when assessing the quality of genome-edited clones, the possibility of the insertion of unexpectedly long exogenous sequences that exceed the detection limit of PCR should always be considered. In this respect, PCR analysis using heterozygous SNPs as shown in Fig. 5 would be useful. However, heterozygous SNPs are not always found in the vicinity of the target site. In such cases, it would be useful to construct two targeting vectors with different selection markers and modify each allele with a different vector.

We performed long-read sequencing with Nanopore technology to obtain a complete picture of the unexpected recombination patterns. Long-read assembly is more effective than short-read assembly when repeat sequences are present. Our analysis detected cases in which two copies of the vectors were linked and inserted into the genome (clone 37 in Fig. 4) or two copies of exon 2 were present after recombination (clones 21, 37, 44, and 45 in Fig. 4). When the backbone of the targeting vector was inserted (clones 6, 24, 31, and 44 in Fig. 4), there was an approximately 3-kb sequence overlap between the cloned insert and the backbone of the cloning vectors. These recombination events would have been difficult to analyze with short-read sequencing. On the other hand, long-read sequencing has the shortcoming of low sequence accuracy. To complement this weakness, we performed direct sequencing of the PCR products shown in Fig. 2C and confirmed the nucleotide sequences of the recombination junctions that were presumed from the long-read sequences. The results of Sanger sequencing matched the consensus sequence inferred from long-read sequencing in almost all cases. Thus, long-read sequencing analysis is a powerful strategy for analyzing complex recombination patterns in which repeat sequences are assumed to exist.

This study showed that MMEJ, NHEJ, and HR are involved in various combinations during the recombination of the vector and the genome. Previous reports have shown that NHEJ occurs throughout the cell cycle, whereas MMEJ and HR are active mainly from the M to the early S phase and from the late S to the G2 phase, respectively (10,14). Thus, we thought that HR and MMEJ were unlikely to coexist. However, MMEJ and HR coexisted in two recombinants analyzed by long-read sequencing (clones 27 and 37 in Fig. 4). This suggests that the range of cell cycle phases with high MMEJ and HR activity could be wider than previously thought, at least in ESCs. Alternatively, it is possible that one recombination event occurred, followed by another after the cell cycle had progressed. In the example in which two vectors were linked and inserted into the genome, MMEJ, NHEJ, and HR were all involved (clone 37 in Figs. 3B, 3C, 4F). This result indicates that different pathways are utilized flexibly to repair double-strand breaks in the genome, which may be a general strategy for organisms to maintain genome integrity.

To efficiently obtain the desired recombinant, it may be worthwhile to increase the activity of the intended recombination pathway or inhibit other recombination pathways. For the PITCh method, it may be useful to increase the efficiency of MMEJ through the forced expression of an MMEJ enhancer such as exonuclease 1 (23) or by using inhibitors of HR or NHEJ (26). To efficiently induce MMEJ, all gRNA target sites should be cleaved simultaneously to expose microhomologies for recombination. For this purpose, using the same gRNA to cut the targeting vector and the genome was proposed previously (24). Alternatively, an improved version of Cas9 (27) or gRNA (28) may be effective. To exclude clones with insertion of the targeting vector backbone into the genome, placing a negative selection marker such as the herpes simplex virus thymidine kinase gene in the backbone and performing negative selection using gancyclovir or 1-2′-deoxy-2′-fluoro-beta-D-arabinofuranosyl-5-iodouracil (FIAU) may be helpful (29).

Taken together, the characterization of unintended recombinants conducted in this study will provide clues for improving the efficiency of genome editing and quality control of genome-edited clones.

## DATA AVAILABILITY

The long-read sequencing data are deposited under the BioProject accession number PRJDB15035.

## ACKNOWLEDGMENTS

We acknowledge the NGS core facility of the Genome Information Research Center at the Research Institute for Microbial Diseases of Osaka University for support with Nanopore sequencing and data analysis. We thank Dr. Junji Takeda for providing KY1.1 ESCs and Dr. Kosuke Yusa for the plasmid pPGKneo-F2F.

