## Supplementary Figures and Table for "Long-read sequence analysis of MMEJ-mediated CRISPR genome editing reveals complex on-target vector insertions that may escape standard PCR-based quality control"

**Supplementary Figure 1. Nucleotide sequence of the targeting vector pKlf-cKO-PITCh.**

1 TCGCGCGTTT CGGTGATGAC GGTGAAAACC TCTGACACAT GCAGCTCCCG GAGACGGTCA CAGCTTGTCT GTAAGCGGAT GCCGGGAGCA GACAAGCCCG

101 TCAGGGCGCG TCAGCGGGTG TTGGCGGGTG TCGGGGCTGG CTTAACTATG CGGCATCAGA GCAGATTGTA CTGAGAGTGC ACCATATGCG GTGTGAAATA

201 CCGCACAGAT GCGTAAGGAG AAAATACCGC ATCAGGCGCC ATTCGCCATT CAGGCTGCGC AACTGTTGGG AAGGGCGATC GGTGCGGGCC TCTTCGCTAT

301 TACGCCAGCT GGCGAAAGGG GGATGTGCTG CAAGGCGATT AAGTTGGGTA ACGCCAGGGT TTTCCCAGTC ACGACGTTGT AAAACGACGG CCAGTGAATT

401 CGAGCTCGGT ACCTCGCGAA TGCATCTAGA TGCATCGTAC GCGTACGTGT TTGGCCAGTA CTGGAGATGA CAAGGTCCAG GGTGCTGACC GCCTATAACT

gRNA-PITCh 5′ microhomology

501 TCGTATAGCA TACATTATAC GAAGTTATCA GGGAGTTAGA CTTCAGGCTG TGGGACAGGA GGTGGGTGCA GGGACTGAGG ACACGCGCGC TGAAGGGATG

loxP

601 CCGTGCACCG GGTGCAGATC TTGAGGGCCT AGTTGTTAGA CTTTGGGGTG CAGGGTAGCA GGAGGCACCC CCACTCACGT CCCGCGCCCT GTCTCCTGCA

701 GCGCTGGCCG CGAAATGAAC CCGAGGCGGG CGGCACGGAT GAGGACCTAA ACAACGTGTT GGACTTCATC CTCTCCATGG GATTGGACGG TCTGGGCGCC

Exon 2 →

801 GAAAATCCTC CCGAGCCCCC GCCGCAGCCC CCGCCGCCTG CCTTCTACTA CCCGGAGCCG GGTGCTCCGC CGCCCTACAG CATCCCCGCG GCCAGCCTGG

901 GAACAGAGCT GCTGCGCCCC GACCTGGACC CGCCTCAGGG GCCGGCTCTG CACGGCCGCT TCCTCCTCGC GCCTCCCGGG CGGCTAGTGA AGGCCGAGCC

1001 CCCCGAGGTG GACGGCGGCG GCTACGGCTG CGCTCCGGGC CTGGCCCACG GACCGCGCGG GCTGAAGCTC GAGGGCGCCC CAGGAGCGAC AGGTGCATGC

1101 ATGCGGGGTC CCGCCGGCCG CCCCCCGCCG CCCCCGGACA CGCCGCCGCT CAGCCCCGAC GGCCCCCTGC GCATCCCGGC GTCCGGTCCC CGCAACCCGT

1201 TCCCGCCGCC CTTCGGTCCC GGCCCCAGCT TCGGCGGTCC CGGCCCCGCG TTGCACTACG GGCCTCCCGC GCCTGGCGCC TTCGGTCTTT TCGAGGACGC

1301 GGCGGCAGCG GCGGCGGCGC TGGGCTTGGC TCCACCTGCC ACGCGCGGTC TCCTCACGCC GCCCTCGTCC CCGCTGGAGC TGCTGGAGGC CAAGCCCAAA

1401 CGCGGCCGCC GCTCCTGGCC CCGCAAGCGC GCCGCCACAC ATACTTGCAG CTACACCAAC TGCGGCAAGA CCTACACCAA GAGCTCGCAC CTAAAGGCGC

1501 ATCTGCGTAC ACACACAGGT GGGCGCCTGG CCTCATTCCC GGGATCTGCG GCAGGGGGAT GGCCGCGAGT TCAGGAACAG GCTAGGTTAG ATATCGCGGC

#### Supplementary Figure 1 (continued)

1601 CGCTCTAGAA CTCCACCGCA TTTAAATGCG GTGGCGGCCG AAGTTCCTAT TCTCTAGAAA GTATAGGAAC TTCGTCGAAG CTAGTGGATC GATCCGAACA

FRT

1701 AACGACCCAA CACCCGTGCG TTTTATTCTG TCTTTTTATT GCCGATCCCC TCAGAAGAAC TCGTCAAGAA GGCGATAGAA GGCGATGCGC TGCGAATCGG

pA ← neo ←

1801 GAGCGGCGAT ACCGTAAAGC ACGAGGAAGC GGTCAGCCCA TTCGCCGCCA AGCTCTTCAG CAATATCACG GGTAGCCAAC GCTATGTCCT GATAGCGGTC

1901 CGCCACACCC AGCCGGCCAC AGTCGATGAA TCCAGAAAAG CGGCCATTTT CCACCATGAT ATTCGGCAAG CAGGCATCGC CATGGGTCAC GACGAGATCC

2001 TCGCCGTCGG GCATGCGCGC CTTGAGCCTG GCGAACAGTT CGGCTGGCGC GAGCCCCTGA TGCTCTTCGT CCAGATCATC CTGATCGACA AGACCGGCTT

2101 CCATCCGAGT ACGTGCTCGC TCGATGCGAT GTTTCGCTTG GTGGTCGAAT GGGCAGGTAG CCGGATCAAG CGTATGCAGC CGCCGCATTG CATCAGCCAT

2201 GATGGATACT TTCTCGGCAG GAGCAAGGTG AGATGACAGG AGATCCTGCC CCGGCACTTC GCCCAATAGC AGCCAGTCCC TTCCCGCTTC AGTGACAACG

2301 TCGAGCACAG CTGCGCAAGG AACGCCCGTC GTGGCCAGCC ACGATAGCCG CGCTGCCTCG TCCTGCAGTT CATTCAGGGC ACCGGACAGG TCGGTCTTGA

2401 CAAAAAGAAC CGGGCGCCCC TGCGCTGACA GCCGGAACAC GGCGGCATCA GAGCAGCCGA TTGTCTGTTG TGCCCAGTCA TAGCCGAATA GCCTCTCCAC

2501 CCAAGCGGCC GGAGAACCTG CGTGCAATCC ATCTTGTTCA ATGGCCGATC CCATATTGGC TGCAGGTCGA AAGGCCCGGA GATGAGGAAG AGGAGAACAG

PGK promoter ←

2601 CGCGGCAGAC GTGCGCTTTT GAAGCGTGCA GAATGCCGGG CCTCCGGAGG ACCTTCGGGC GCCCGCCCCG CCCCTGAGCC CGCCCCTGAG CCCGCCCCCG

2701 GACCCACCCC TTCCCAGCCT CTGAGCCCAG AAAGCGAAGG AGCAAAGCTG CTATTGGCCG CTGCCCCAAA GGCCTACCCG CTTCCATTGC TCAGCGGTGC

2801 TGTCCATCTG CACGAGACTA GTGAGACGTG CTACTTCCAT TTGTCACGTC CTGCACGACG CGAGCTGCGG GGCGGGGGGG AACTTCCTGA CTAGGGGAGG

2901 AGTAGAAGGT GGCGCGAAGG GGCCACCAAA GAACGGAGCC GGTTGGCGCC TACCGGTGGA TGTGGAATGT GTGCGAGGCC AGAGGCCACT TGTGTAGCGC

3001 CAAGTGCCCA GCGGGGCTGC TAAAGCGCAT GCTCCAGACT GCCTTGGGAA AAGCGCCTCC CCTACCCGGT AGAATTGACC TGCAGGGGCC CTCGAGAAGT

#### Supplementary Figure 1 (continued)

3101 TCCTATTCTC TAGAAAGTAT AGGAACTTCA TAACTTCGTA TAGCATACAT TATACGAAGT TATGGCGCGC CCAGAGGAAG ACCCTGTCCA TCCTCCAGAA

FRT loxP 3′ microhomology

3201 GGGAAATGGA TCCAAACACG TACGCGTACG ATGCATCGGA TCCCGGGCCC GTCGACTGCA GAGGCCTGCA TGCAAGCTTG GCGTAATCAT GGTCATAGCT

gRNA-PITCh

3301 GTTTCCTGTG TGAAATTGTT ATCCGCTCAC AATTCCACAC AACATACGAG CCGGAAGCAT AAAGTGTAAA GCCTGGGGTG CCTAATGAGT GAGCTAACTC

3401 ACATTAATTG CGTTGCGCTC ACTGCCCGCT TTCCAGTCGG GAAACCTGTC GTGCCAGCTG CATTAATGAA TCGGCCAACG CGCGGGGAGA GGCGGTTTGC

3501 GTATTGGGCG CTCTTCCGCT TCCTCGCTCA CTGACTCGCT GCGCTCGGTC GTTCGGCTGC GGCGAGCGGT ATCAGCTCAC TCAAAGGCGG TAATACGGTT

3601 ATCCACAGAA TCAGGGGATA ACGCAGGAAA GAACATGTGA GCAAAAGGCC AGCAAAAGGC CAGGAACCGT AAAAAGGCCG CGTTGCTGGC GTTTTTCCAT

3701 AGGCTCCGCC CCCCTGACGA GCATCACAAA AATCGACGCT CAAGTCAGAG GTGGCGAAAC CCGACAGGAC TATAAAGATA CCAGGCGTTT CCCCCTGGAA

3801 GCTCCCTCGT GCGCTCTCCT GTTCCGACCC TGCCGCTTAC CGGATACCTG TCCGCCTTTC TCCCTTCGGG AAGCGTGGCG CTTTCTCATA GCTCACGCTG

3901 TAGGTATCTC AGTTCGGTGT AGGTCGTTCG CTCCAAGCTG GGCTGTGTGC ACGAACCCCC CGTTCAGCCC GACCGCTGCG CCTTATCCGG TAACTATCGT

4001 CTTGAGTCCA ACCCGGTAAG ACACGACTTA TCGCCACTGG CAGCAGCCAC TGGTAACAGG ATTAGCAGAG CGAGGTATGT AGGCGGTGCT ACAGAGTTCT

4101 TGAAGTGGTG GCCTAACTAC GGCTACACTA GAAGAACAGT ATTTGGTATC TGCGCTCTGC TGAAGCCAGT TACCTTCGGA AAAAGAGTTG GTAGCTCTTG

4201 ATCCGGCAAA CAAACCACCG CTGGTAGCGG TGGTTTTTTT GTTTGCAAGC AGCAGATTAC GCGCAGAAAA AAAGGATCTC AAGAAGATCC TTTGATCTTT

4301 TCTACGGGGT CTGACGCTCA GTGGAACGAA AACTCACGTT AAGGGATTTT GGTCATGAGA TTATCAAAAA GGATCTTCAC CTAGATCCTT TTAAATTAAA

4401 AATGAAGTTT TAAATCAATC TAAAGTATAT ATGAGTAAAC TTGGTCTGAC AGTTACCAAT GCTTAATCAG TGAGGCACCT ATCTCAGCGA TCTGTCTATT

Amp ←

4501 TCGTTCATCC ATAGTTGCCT GACTCCCCGT CGTGTAGATA ACTACGATAC GGGAGGGCTT ACCATCTGGC CCCAGTGCTG CAATGATACC GCGAGACCCA

4601 CGCTCACCGG CTCCAGATTT ATCAGCAATA AACCAGCCAG CCGGAAGGGC CGAGCGCAGA AGTGGTCCTG CAACTTTATC CGCCTCCATC CAGTCTATTA

4701 ATTGTTGCCG GGAAGCTAGA GTAAGTAGTT CGCCAGTTAA TAGTTTGCGC AACGTTGTTG CCATTGCTAC AGGCATCGTG GTGTCACGCT CGTCGTTTGG

4801 TATGGCTTCA TTCAGCTCCG GTTCCCAACG ATCAAGGCGA GTTACATGAT CCCCCATGTT GTGCAAAAAA GCGGTTAGCT CCTTCGGTCC TCCGATCGTT

#### Supplementary Figure 1 (continued)

4901 GTCAGAAGTA AGTTGGCCGC AGTGTTATCA CTCATGGTTA TGGCAGCACT GCATAATTCT CTTACTGTCA TGCCATCCGT AAGATGCTTT TCTGTGACTG

5001 GTGAGTACTC AACCAAGTCA TTCTGAGAAT AGTGTATGCG GCGACCGAGT TGCTCTTGCC CGGCGTCAAT ACGGGATAAT ACCGCGCCAC ATAGCAGAAC

5101 TTTAAAAGTG CTCATCATTG GAAAACGTTC TTCGGGGCGA AAACTCTCAA GGATCTTACC GCTGTTGAGA TCCAGTTCGA TGTAACCCAC TCGTGCACCC

5201 AACTGATCTT CAGCATCTTT TACTTTCACC AGCGTTTCTG GGTGAGCAAA AACAGGAAGG CAAAATGCCG CAAAAAAGGG AATAAGGGCG ACACGGAAAT

5301 GTTGAATACT CATACTCTTC CTTTTTCAAT ATTATTGAAG CATTTATCAG GGTTATTGTC TCATGAGCGG ATACATATTT GAATGTATTT AGAAAAATAA

5401 ACAAATAGGG GTTCCGCGCA CATTTCCCCG AAAAGTGCCA CCTGACGTCT AAGAAACCAT TATTATCATG ACATTAACCT ATAAAAATAG GCGTATCACG

5501 AGGCCCTTTC GTC

#### Supplementary Figure 2. Consensus sequence of Clone 37 obtained using Flye-assembler.

Nanopore long sequences are prone to errors in regions where the same nucleotides are repeated. Differences between the Nanopore sequencing data and the C57BL/6J mouse genome reference sequence are indicated by boxing the nucleotides and showing the corresponding reference sequence. The regions confirmed by Sanger sequencing (Figs. 3B, 3C) are also indicated below.

GCTCAGCGAGCCTATCTTGCCGTCCTTTGCCACTTTCGCCAGCCCGTGCGAGCGCGGCCTCCAGGAGGTGAGGGCAGCGAGGGTCCGCGGGCGG

primer

Confirmed by Sanger sequencing (Region I in Figs. 3B, 3C)

CGGGGTAGCAAGGTACCAGGCTACAGGGTGCAGGAGCCAGTACTGGAGATGACAAGGTCCAGGGTGCTGACCTCCCAGGGAGTTAGACTTCAGG

5′ microhomology (4-bp deletion) 3-bp insertion gRNA-1 target (3′ region downstream of the cleavage site)

CTGTGGGACAGGAGGTGGGTGCAGGGACTGAGGACACGCGCGCTGAAGGGATGCCGTGCACCGGGTGCAGATCTTGAGGGCCTAGTTGTTAGAC

Exon 2

TTTGGGGTGCAGGGTAGCAGGAGGCACCCCCACTCACGTCCCGCGCCCTGTCTCCTGCAGCGCTGGCCGCGAAATGAACCCGAGGCGGGCGGCA

Ref seq: CCCCC

CGGATGAGGACCTAAACAACGTGTTGGACTTCATCCTCTCCATGGGATTGGACGGTCTGGGCGCCGAAAATCCTCCCGAGCCCCGCCGCAGCCC

CCGCCGCCTGCCTTCTACTACCCGGAGCCGGGTGCTCCGCCGCCCTACAGCATCCCCGCGGCCAGCCTGGGAACAGAGCTGCTGCGCCCCGACC

Ref seq: CCCCCC

TGGACCCGCCTCAGGGGCCGGCTCTGCACGGCCGCTTCCTCCTCGCGCCTCCCGGGCGGCTAGTGAAGGCCGAGCCCCCGAGGTGGACGGCGGC

GGCTACGGCTGCGCTCCGGGCCTGGCCCACGGACCGCGCGGGCTGAAGCTCGAGGGCGCCCCAGGAGCGACAGGTGCATGCATGCGGGGTCCCG

Ref seq: CCCCCC Ref seq: CCCCC

CCGGCCGCCCCCGCCGCCCCGGACACGCCGCCGCTCAGCCCCGACGGCCCCCTGCGCATCCCGGCGTCCGGTCCCCGCAACCCGTTCCCGCCGC

CCTTCGGTCCCGGCCCCAGCTTCGGCGGTCCCGGCCCCGCGTTGCACTACGGGCCTCCCGCGCCTGGCGCCTTCGGTCTTTTCGAGGACGCGGC

GGCAGCGGCGGCGGCGCTGGGCTTGGCTCCACCTGCCACGCGCGGTCTCCTCACGCCGCCCTCGTCCCCGCTGGAGCTGCTGGAGGCCAAGCCC

AAACGCGGCCGCCGCTCCTGGCCCCGCAAGCGCGCCGCCACACATACTTGCAGCTACACCAACTGCGGCAAGACCTACACCAAGAGCTCGCACC

Exon 2 Ref seq: CCC Ref seq: GGGGG

TAAAGGCGCATCTGCGTACACACACAGGTGGGCGCCTGGCCTCATTCCGGGATCTGCGGCAGGGGATGGCCGCGAGTTCAGGAACAGGCTAGGT

#### Supplementary Figure 2 (continued)

TAGATATCGCGGCCGCTCTAGAACTCCACCGCATTTAAATGCGGTGGCGGCCGAAGTTCCTATTCTCTAGAAAGTATAGGAACTTCGTCGAAGC

FRT

Ref seq: CCCC

TAGTGGATCGATCCGAACAAACGACCCAACACCCGTGCGTTTTATTCTGTCTTTTTATTGCCGATCCCTCAGAAGAACTCGTCAAGAAGGCGAT

PGKneopA (1,222-bp deletion)

AGAAGGCGATGCGCTGCGAATCGGGAGCGGCGATACCGTAAAGCACGAGGAAGCGGTCAGCCCATTCGCCGCCAAGCTCTTCAGCAATATCACG

GGTAGCCAACGCTATGTCCTGATAGCGGTCCGCCACACCCAGCCGGCCACAGTCGATGAATCCAGAAAAGCGGCCATTTTCCACCATGATATTC

Confirmed by Sanger sequencing (Region II in Figs. 3B, 3C)

GGCAAGCAGGCATCGCCATGGGTCACGACGATGGGTCAAACTGGTGTTTGGCCAGTACTGGAGATGACAAGGTCCAGGGTGCTGACCGCCTATA

14-bp insertion

5′ microhomology

ACTTCGTATAGCATACATTATACGAAGTTATCAGGGAGTTAGACTTCAGGCTGTGGGACAGGAGGTGGGTGCAGGGACTGAGGACACGCGCGCT

loxP

Ref seq: T

Ref seq: CCCCC

GAAGGGATGCCGTGCACCGGGTGCAGATCTTGAGGGCCTAGTTGTTAGACTTTGGGGCGCAGGGTAGCAGGAGGCACCCCACTCACGTCCCGCG

Exon 2

CCCTGTCTCCTGCAGCGCTGGCCGCGAAATGAACCCGAGGCGGGCGGCACGGATGAGGACCTAAACAACGTGTTGGACTTCATCCTCTCCATGG

GATTGGACGGTCTGGGCGCCGAAAATCCTCCCGAGCCCCCGCCGCAGCCCCCGCCGCCTGCCTTCTACTACCCGGAGCCGGGTGCTCCGCCGCC

Ref seq: GC

CTACAGCATCCCCGCGGCCAGCCTGGGAACAGAGCTGCTGCACCCGACCTGGACCCGCCTCAGGGGCCGGCTCTGCACGGCCGCTTCCTCCTCG

Ref seq: CCCCCC

CGCCTCCCGGGCGGCTAGTGAAGGCCGAGCCCCCGAGGTGGACGGCGGCGGCTACGGCTGCGCTCCGGGCCTGGCCCACGGACCGCGCGGGCTG

Ref seq: CCCCCC

Ref seq: CCCCC

AAGCTCGAGGGCGCCCCAGGAGCGACAGGTGCATGCATGCGGGGTCCCGCCGGCCGCCCCCGCCGCCCCGGACACGCCGCCGCTCAGCCCCGAC

GGCCCCCTGCGCATCCCGGCGTCCGGTCCCCGCAACCCGTTCCCGCCGCCCTTCGGTCCCGGCCCCAGCTTCGGCGGTCCCGGCCCCGCGTTGC

ACTACGGGCCTCCCGCGCCTGGCGCCTTCGGTCTTTTCGAGGACGCGGCGGCAGCGGCGGCGGCGCTGGGCTTGGCTCCACCTGCCACGCGCGG

### Supplementary Figure 2 (continued)

TCTCCTCACGCCGCCCTCGTCCCCGCTGGAGCTGCTGGAGGCCAAGCCCAAACGCGGCCGCCGCTCCTGGCCCCGCAAGCGCGCCGCCACACAT

Exon 2

ACTTGCAGCTACACCAACTGCGGCAAGACCTACACCAAGAGCTCGCACCTAAAGGCGCATCTGCGTACACACACAGGTGGGCGCCTGGCCTCAT

Ref seq: CCC    Ref seq: GGGGG

TCCGGGATCTGCGGCAGGGGATGGCCGCGAGTTCAGGAACAGGCTAGGTTAGATATCGCGGCCGCTCTAGAACTCCACCGCATTTAAATGCGGT

GGCGGCCGAAGTTCCTATTCTCTAGAAAGTATAGGAACTTCGTCGAAGCTAGTGGATCGATCCGAACAAACGACCCAACACCCGTGCGTTTTAT

FRT

PGKneopA

Ref seq: CCCC

TCTGTCTTTTTATTGCCGATCCCTCAGAAGAACTCGTCAAGAAGGCGATAGAAGGCGATGCGCTGCGAATCGGGAGCGGCGATACCGTAAAGCA

CGAGGAAGCGGTCAGCCCATTCGCCGCCAAGCTCTTCAGCAATATCACGGGTAGCCAACGCTATGTCCTGATAGCGGTCCGCCACACCCAGCCG

GCCACAGTCGATGAATCCAGAAAAGCGGCCATTTTCCACCATGATATTCGGCAAGCAGGCATCGCCATGGGTCACGACGAGATCCTCGCCGTCG

GGCATGCGCGCCTTGAGCCTGGCGAACAGTTCGGCTGGCGCGAGCCCCTGATGCTCTTCGTCCAGATCATCCTGATCGACAAGACCGGCTTCCA

TCCGAGTACGTGCTCGCTCGATGCGATGTTTCGCTTGGTGGTCGAATGGGCAGGTAGCCGGATCAAGCGTATGCAGCCGCCGCATTGCATCAGC

CATGATGGATACTTTCTCGGCAGGAGCAAGGTGAGATGACAGGAGATCCTGCCCCGGCACTTCGCCCAATAGCAGCCAGTCCCTTCCCGCTTCA

GTGACAACGTCGAGCACAGCTGCGCAAGGAACGCCCGTCGTGGCCAGCCACGATAGCCGCGCTGCCTCGTCCTGCAGTTCATTCAGGGCACCGG

Ref seq: AAAAA

ACAGGTCGGTCTTGACAAAAGAACCGGGCGCCCCTGCGCTGACAGCCGGAACACGGCGGCATCAGAGCAGCCGATTGTCTGTTGTGCCCAGTCA

TAGCCGAATAGCCTCTCCACCCAAGCGGCCGGAGAACCTGCGTGCAATCCATCTTGTTCAATGGCCGATCCCATATTGGCTGCAGGTCGAAAGG

CCCGGAGATGAGGAAGAGGAGAACAGCGCGGCAGACGTGCGCTTTTGAAGCGTGCAGAATGCCGGGCCTCCGGAGGACCTTCGGGCGCCCGCCC

### Supplementary Figure 2 (continued)

Ref seq: CCCCC Ref seq: CCCC

CGCCCCTGAGCCCGCCCCTGAGCCCGCCCCGGACCCACCCTTCCCAGCCTCTGAGCCCAGAAAGCGAAGGAGCAAAGCTGCTATTGGCCGCTGC

CCCAAAGGCCTACCCGCTTCCATTGCTCAGCGGTGCTGTCCATCTGCACGAGACTAGTGAGACGTGCTACTTCCATTTGTCACGTCCTGCACGA

Ref seq: GGGGGGG

CGCGAGCTGCGGGGCGGGGGAACTTCCTGACTAGGGGAGGAGTAGAAGGTGGCGCGAAGGGGCCACCAAAGAACGGAGCCGGTTGGCGCCTACC

GGTGGATGTGGAATGTGTGCGAGGCCAGAGGCCACTTGTGTAGCGCCAAGTGCCCAGCGGGGCTGCTAAAGCGCATGCTCCAGACTGCCTTGGG

Ref seq: CCCC

AAAAGCGCCTCCCTACCCGGTAGAATTGACCTGCAGGGGCCCTCGAGAAGTTCCTATTCTCTAGAAAGTATAGGAACTTCATAACTTCGTATAG

PGKneopA FRT

Confirmed by Sanger sequencing (Region III in Figs. 3B, 3C)

CATACATTATACGAAGTTATGGCGCGCCCAGAGGAAGACCCTGTCCATCCTCCAGAAGGGAAATGGATACTGTCCCCAGAGTCTATGACACTGC

loxP 3′ microhomology

Ref seq: GGGGG Ref seq: GGG

CAAGGGGTGTGGCAGTTTAATGGGTTGGCTCCGAGGACTCAGGTCTCAAGAATTCGGGACATAATTAAGAAGCTGTGGAGGGCTGG

primer

**Supplementary Figure 3. Annotation of consensus sequences obtained by de novo assembly of long-read sequencing data and confirmation of assembly fidelity by Sanger direct sequencing.**

### Clone 6

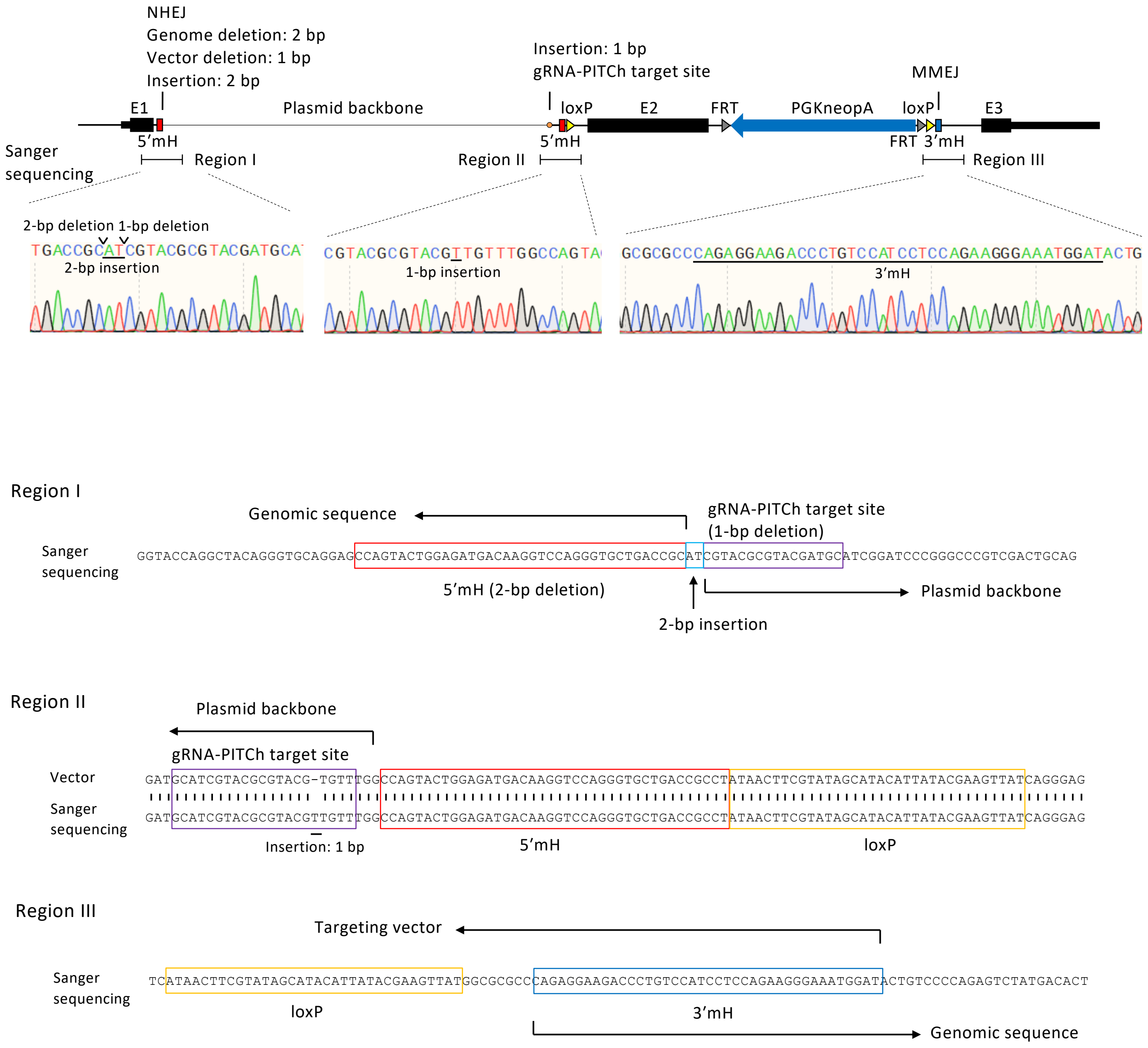

#### Supplementary Figure 3 (continued)

### Clone 21

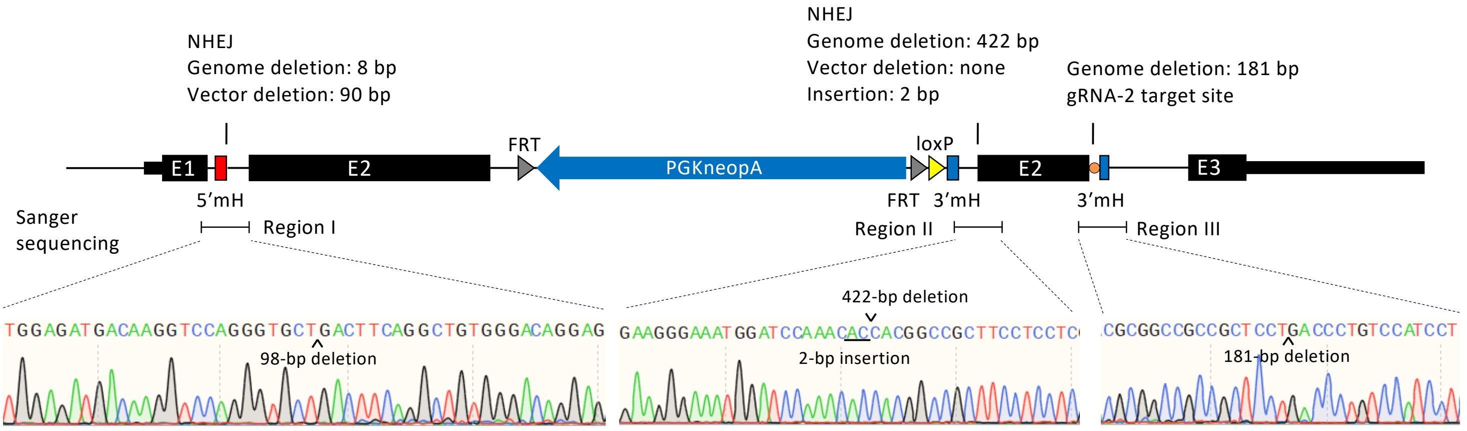


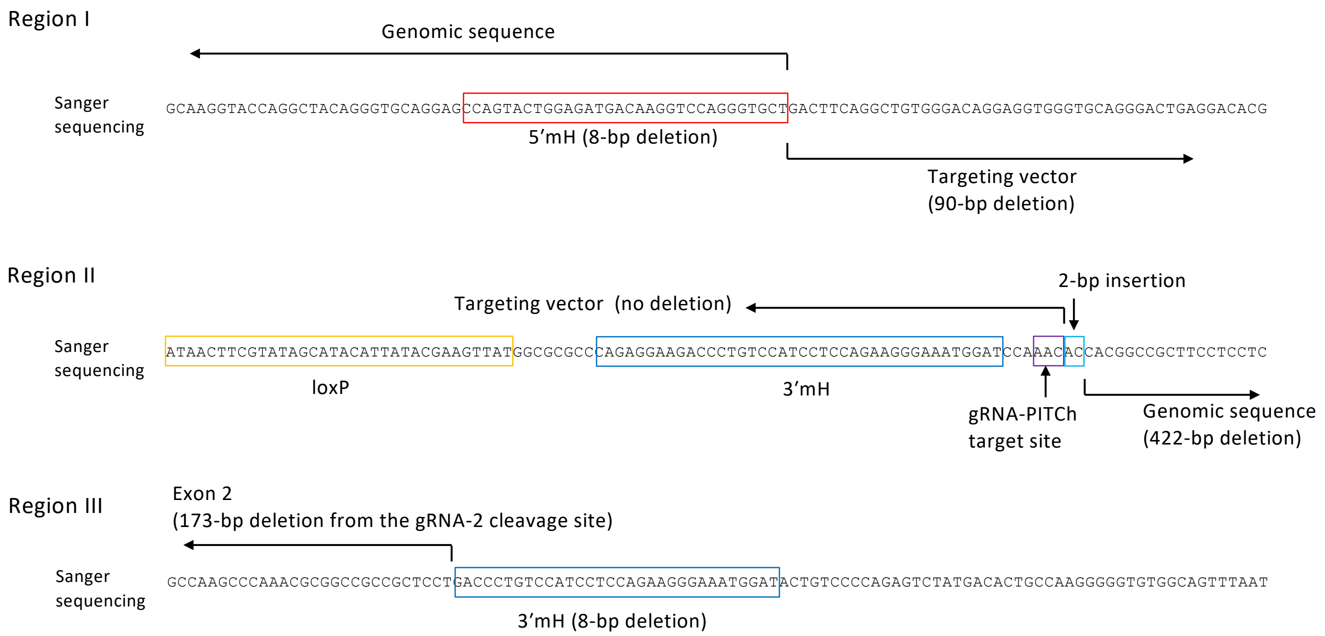

#### Supplementary Figure 3 (continued)

### Clone 24

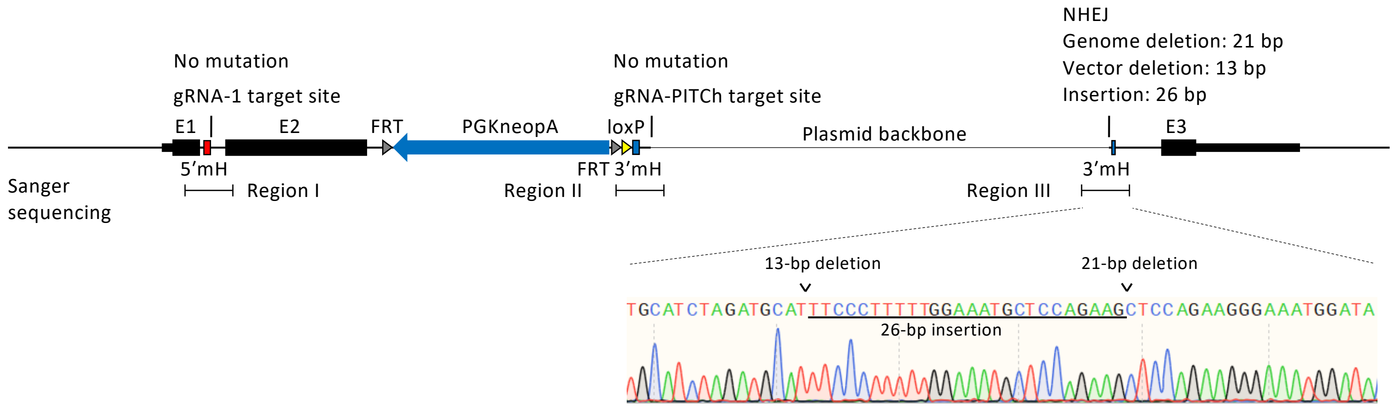


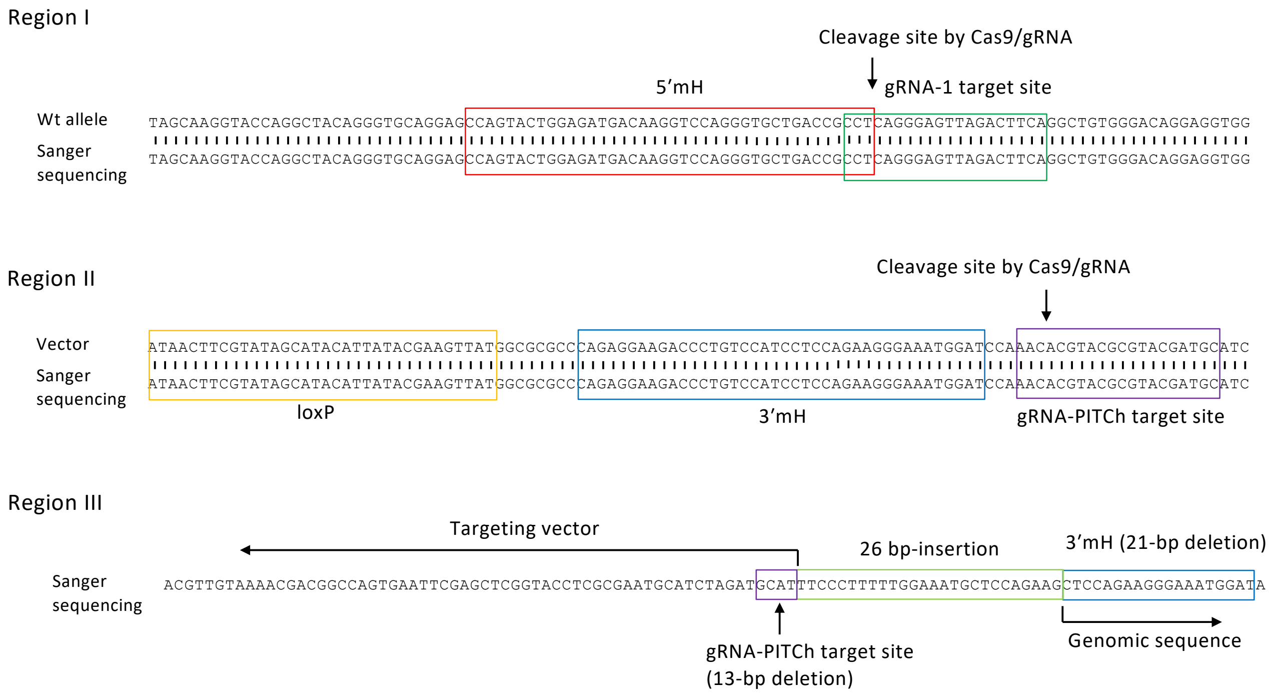

**Supplementary Figure 3 (continued)**

### Clone 27

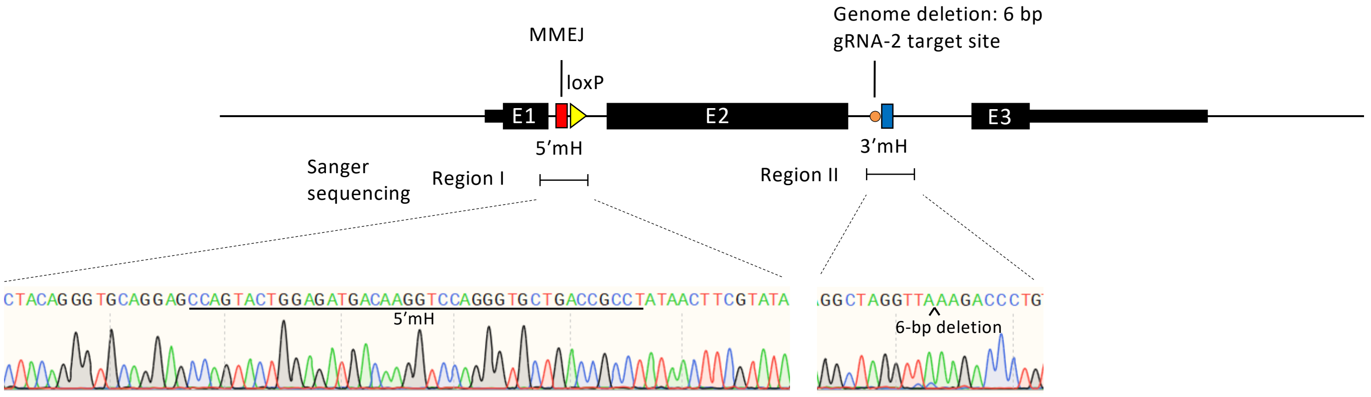


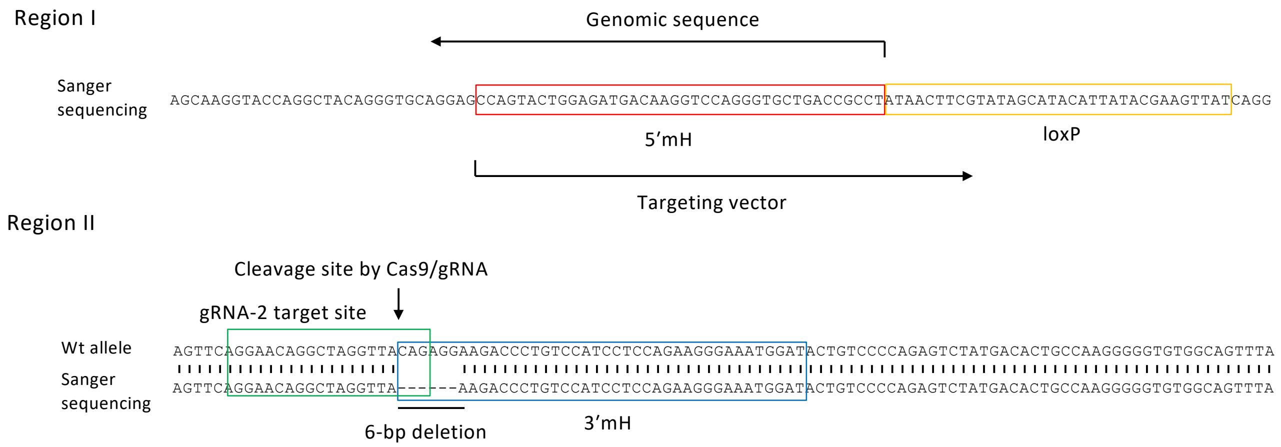

#### Supplementary Figure 3 (continued)

### Clone 29

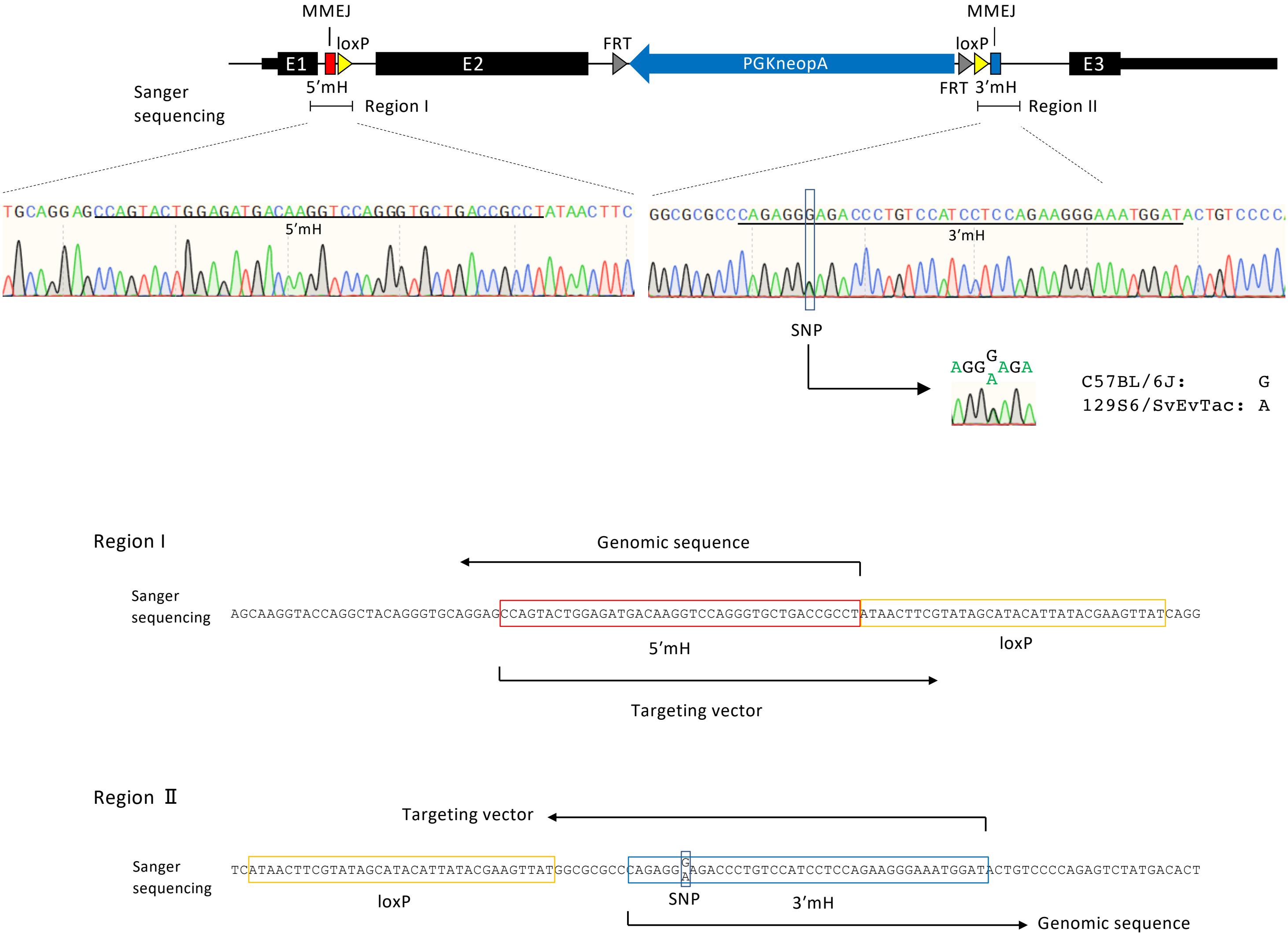

#### Supplementary Figure 3 (continued)

### Clone 31

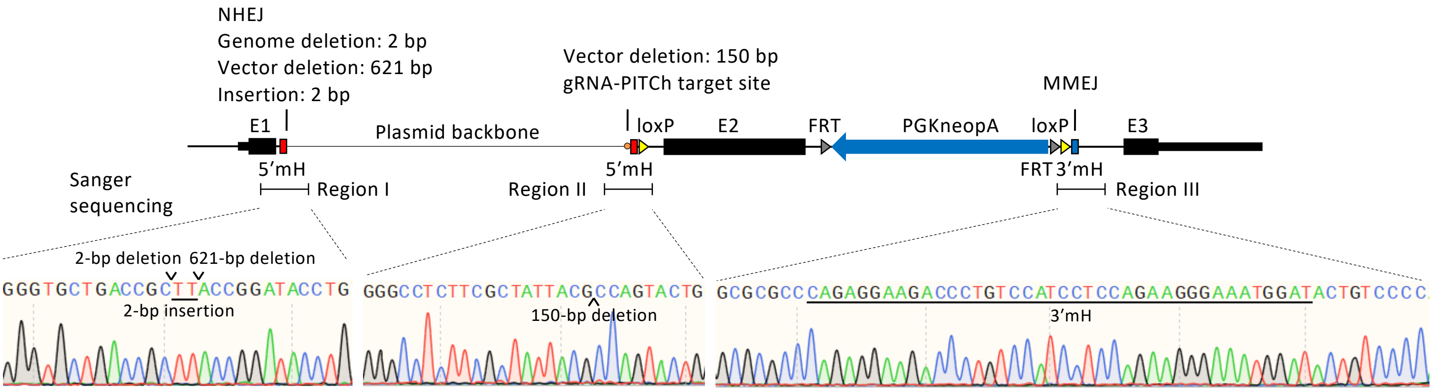


Region I

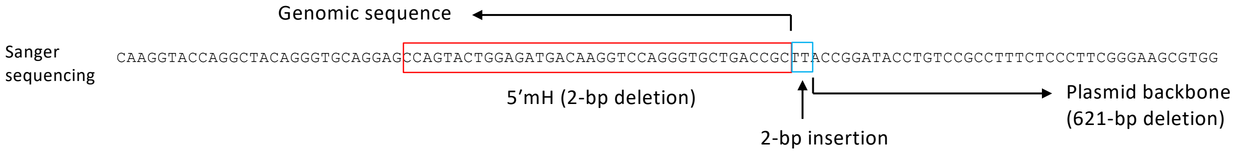


Region Ⅱ

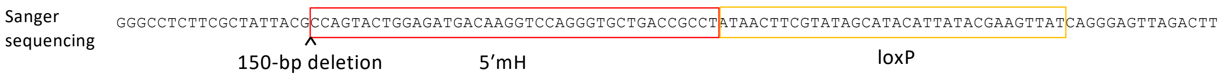


Region III

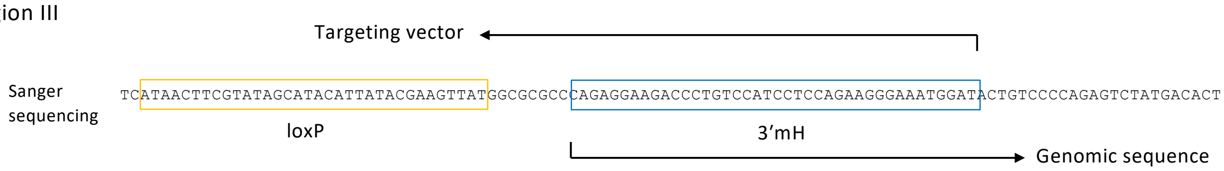

#### Supplementary Figure 3 (continued)

### Clone 39

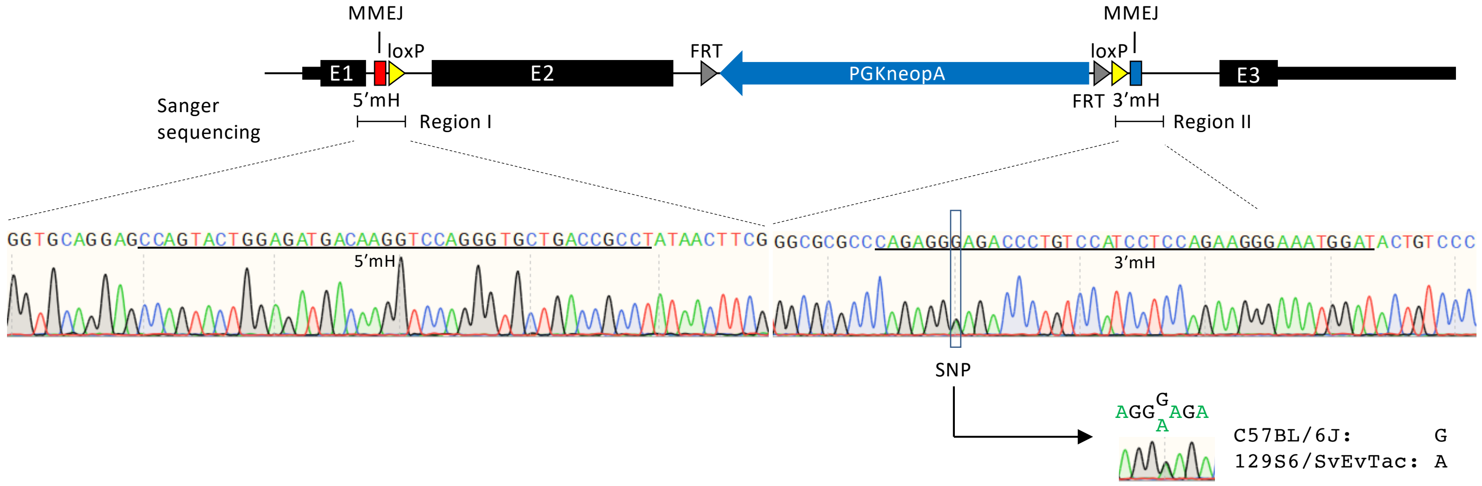


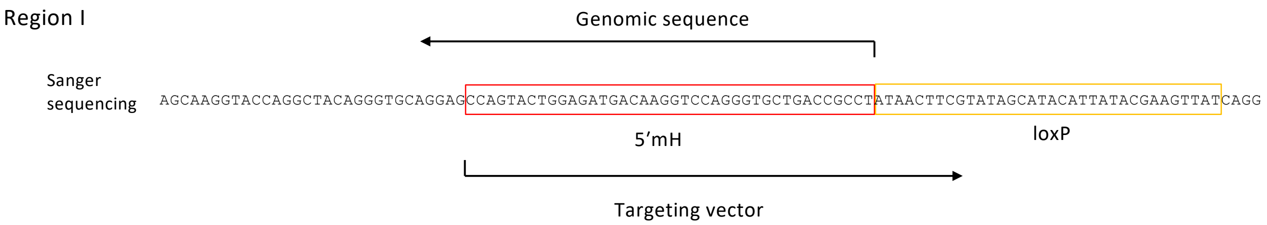


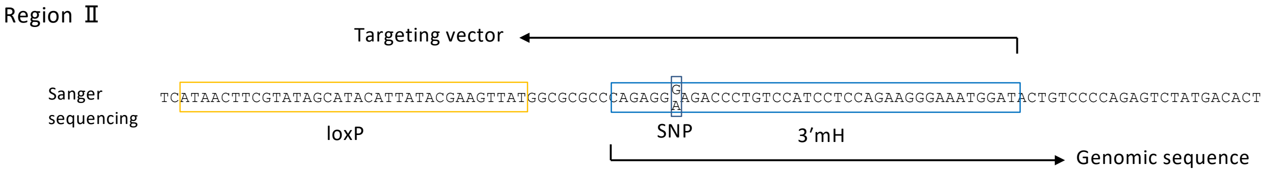

**Supplementary Figure 3 (continued)**

### Clone 44

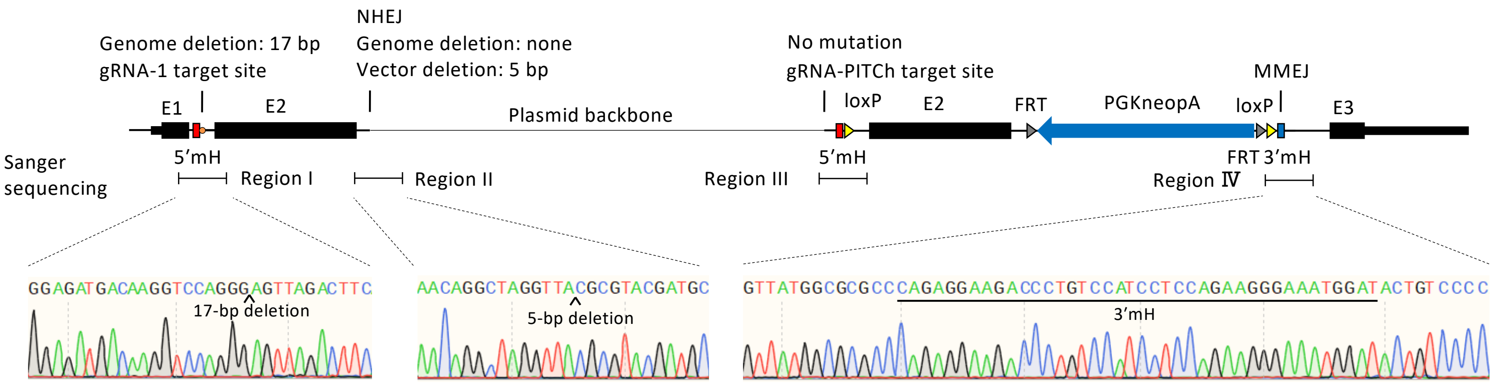


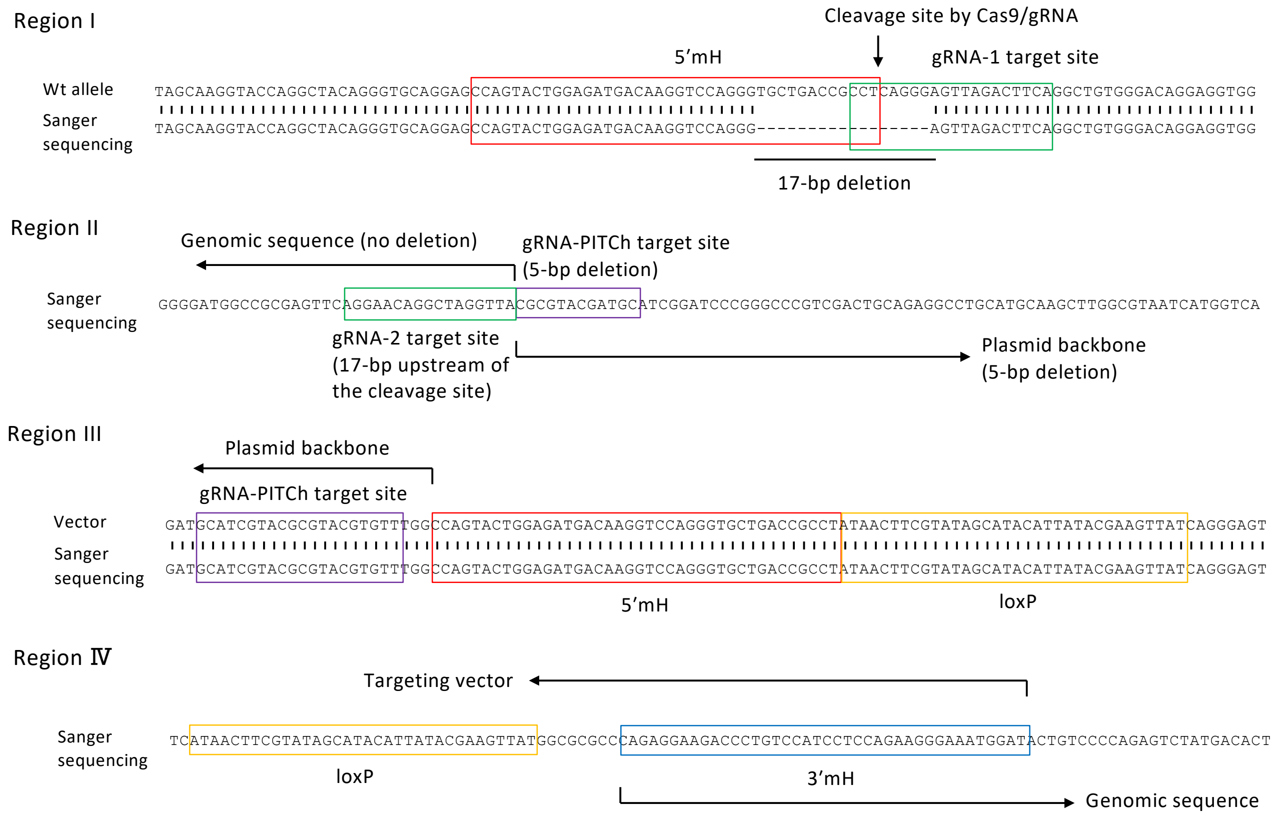

#### Supplementary Figure 3 (continued)

### Clone 45

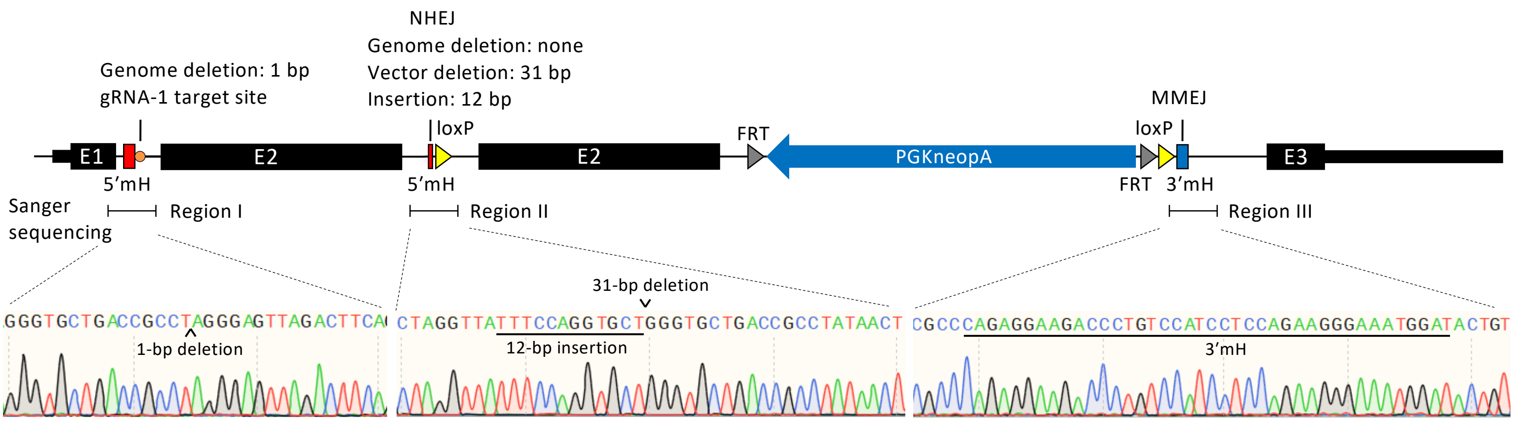


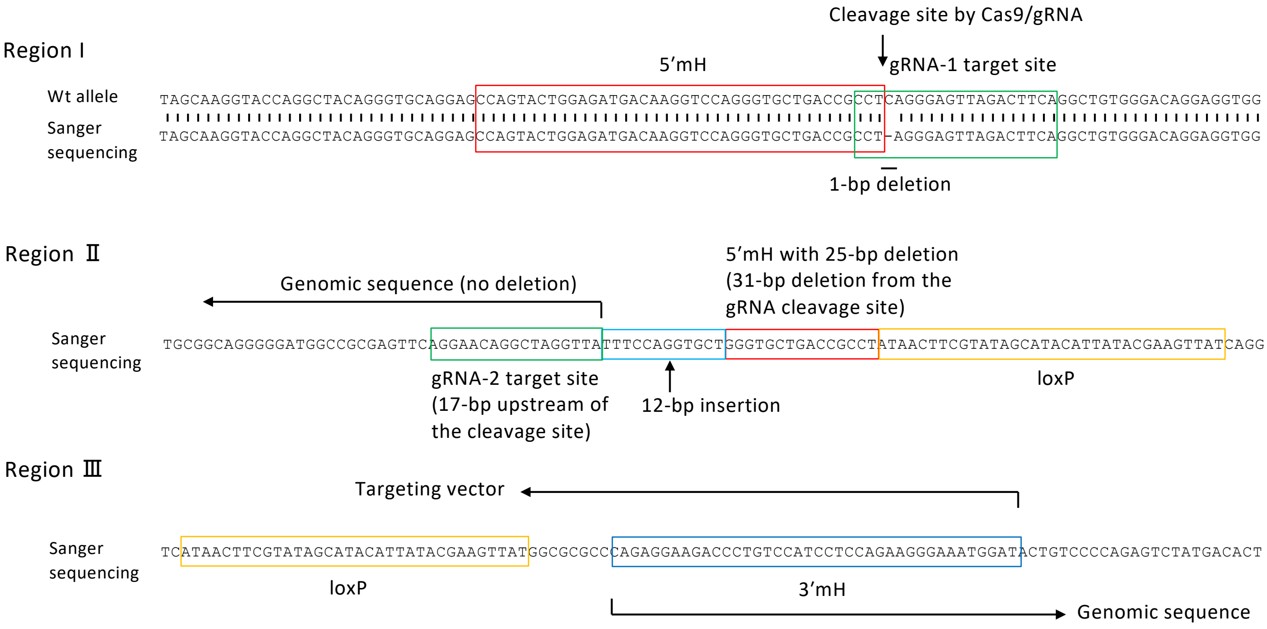

**Supplementary Table 1. Sequences of the oligonucleotides used in this study.**

| | |
|---|---|
| Construction of the Cas9/gRNA vector | |
| pX330-PITCh | |
| gPITCH-F2 | caccGCATCGTACGCGTACGTGTT |
| gPITCH-R2 | aaacAACACGTACGCGTACGATGC |
| pX330-gRNA-1 | |
| gKlf2-4-F | caccgTGAAGTCTAACTCCCTGAGG |
| gKlf2-4-R | aaacCCTCAGGGAGTTAGACTTCAc |
| pX330-gRNA-2 | |
| gKlf2-7-F | caccgAGGAACAGGCTAGGTTACAG |
| gKlf2-7-R | aaacCTGTAACCTAGCCTGTTCCTc |
| Construction of the targeting vector | |
| Klf2-5HR-F1 | ACAAAAGCTGGAGCTCCACGTAGCCAAAGGGGCCTTGAACCTAATG |
| Klf2-5HR-R1 | CGTATAATGTATGCTATACGAAGTTATAGGCGGTCAGCACCCTGGACCTT |
| Klf2-5HR-F2 | CGTATAGCATACATTATACGAAGTTATCAGGGAGTTAGACTTCAGGCTGT |
| Klf2-5HR-R2 | GTGGAGTTCTAGAGCGGCCGCGATATCTAACCTAGCCTGTTCCTGAACTC |
| Detection of upstream recombination (primer I, Fig. 2) | |
| Klf2-5mH-scr1 | GCTCAGCGAGCCTATCTTGCCGTCCTTT |
| Klf2-loxP-F1 | CCAAAGTCTAACAACTAGGCCCTCAAG |
| Detection of the full-length insertion sequence (primer II, Fig. 2) | |
| Klf2-5mH-scr1 | GCTCAGCGAGCCTATCTTGCCGTCCTTT |
| Klf2-3mH-scr2 | CCAGCCCTCCACAGCTTCTTAATTATGTC |
| Detection of downstream recombination (primer III, Fig. 2) | |
| PGK-R1 | CGGGGCTGCTAAAGCGCATGCTCCAGACTG |
| Klf2-3mH-scr2 | CCAGCCCTCCACAGCTTCTTAATTATGTC |
| Detection of SNPs (Fig. 5) | |
| Klf2-SNP-5mH-F1 | CAAGGTAGCTTAAACAAAGATTTCACAGAG |
| Klf2-SNP-3mH-R1 | TGCATCAGAGAGGAATGACATTGAGC |
